# Test-retest reproducibility of human brain multi-slice 1H FID-MRSI data at 9.4 T after optimization of lipid regularization, macromolecular model and spline baseline stiffness

**DOI:** 10.1101/2022.02.02.478678

**Authors:** Theresia Ziegs, Andrew Martin Wright, Anke Henning

## Abstract

**Purpose:** This study analyzes the effects of retrospective lipid suppression, a simulated macromolecular prior knowledge and different spline baseline stiffness values on 9.4 T multi-slice proton FID-MRSI data spanning the whole cerebrum of human brain and its reproducibility of metabolite ratio (/tCr) maps for 10 brain metabolites.

**Methods:** Measurements were performed twice on five volunteers using a non-accelerated FID MRSI 2D sequence at 9.4 T. The effects of retrospective lipid L2-regularization, macromolecular spectrum and different LCModel baseline flexibilities on SNR, FWHM, fitting residual, CRLB and the concentration ratio maps were investigated. Intra-subject, inter-session coefficient of variation of the mean metabolite ratios (/tCr) of each slice was calculated.

**Results:** L2-regularization provided effective suppression of lipid-artifacts, but should be avoided if no artifacts are detected. Transversal, sagittal and coronal of many metabolite ratio maps correspond to anatomically expected concentration relations in gray and white matter for the majority of the cerebrum when using a flexible baseline in LCModel fit. Additionally, results from the second measurements of the same subjects show that slice positioning and data quality correlate significantly to the first measurement.

**Conclusion:** Concentration ratio maps (/tCr) for 4 metabolites (tCho, NAA, Glu, mI) spanning the majority and six metabolites (NAAG, GABA, GSH, Tau, Gln, Asp) covering 32 mm in the upper part of the brain were acquired at 9.4 T using multi-slice FID MRSI with retrospective lipid suppression, a macromolecular spectrum and a flexible LCModel baseline.

## 1. Introduction

Metabolite mapping with ^1^H MRSI spanning a major part of the brain provides a comprehensive insight into the distribution of metabolites across different regions in the human brain. Thus, it is a valuable tool to obtain complimentary information to MRI, diffusion or fMRI techniques regarding brain metabolism^1^, and has been used to examine for various diseases^2–10^.

Most whole-brain 1H MRSI studies have been conducted at field strengths ≤ 3T^11–26^ showing concentration or ratio maps of maximal five metabolites NAA, Cr, Cho, mI and Glx. However, a study from Motyka et al.^27^ compared whole-brain simulations of MRSI data for different field strengths from 1.5T, 3T, 7T up to 9.4T and concluded that MRSI benefits significantly from ultra-high field strengths in terms of SNR and spectral resolution. Ultra-high fields enable the detection of low concentrated metabolites such as N-acetylaspartylglutamate (NAAG), and the distinction between metabolites that are overlapped at lower field strengths (< 3T) such as glutamate (Glu) and glutamine (Gln).

While the majority of ultra-high field ^1^H MRSI studies cover only a single slice^28–35^, there are only very few ^1^H MRSI investigations at ultra-high field strengths that show an extended coverage of the human brain^36–41^. These publications were all using acceleration techniques and showing maps of a maximum of six metabolites (tCr, tNAA/NAA, tCho, Glu, Gln, mI) for a brain volume of a maximum of 110 mm and only one paper showed NAAG for a relatively small volume of 50 mm thickness^39^. None of these ultra-high field ^1^H MRSI studies investigated the intra-subject test-retest reproducibility, only one ultra-high field MRSI study investigated inter-subject reproducibility^37^. In fact there is only one 2D ^1^H MRSI study at 7T that presents intra-subject test-retest data from a single slice above the corpus callosum for tNAA, tCho, tCr, mI and Glx^42^.

In this paper, the number of metabolites quantified by ^1^H MRSI from the human brain shall be extended and concentration ratio (/tCr) maps for 10 brain metabolites will be obtained, namely for NAA, tCho, Glu, and mI for the whole cerebrum and Gln, γ-aminobutyric acid (GABA), glutathione (GSH), NAAG, taurine (Tau) and aspartate (Asp) for a stack of FID-MRSI slices covering 32 mm in the upper part of the human brain. In addition, a test-retest analysis verified inter-session, intra-subject reproducibly of the multi-slice ^1^H FID MRSI measurements of these brain metabolites. For this purpose, the post-processing steps were systematically optimized and validated with optional lipid removal and macromolecular (MM) spectrum included in the fitting routine. Since these both influence the data’s baseline, the impact of the spline baseline flexibility was investigated, which has not been done before for 1H MRSI data, although accurate quantification rely strongly on baseline estimations^43,44^.

## 2. Methods

### 2.1. Data Acquisition

#### 2.1.1 Hardware

All measurements were performed using a 9.4T Magnetom whole-body MR scanner (Siemens Healthineers, Erlangen, Germany) using an in-house built radiofrequency (RF) array coil with 18 transmit and 32 receive channels^45^. The receive channels consist of 18 transceiver surface loops and 14 receive only vertical loops in a double-row design.

#### 2.1.2 Study-design

Five healthy volunteers (4 male and 1 female, aged 28 ± 2 years) provided written informed consent according to the local ethics board regulations and were scanned twice each. Repositioning of each subject was carefully carried out by using the same padding under the head and neck and at the sides inside the tight fit RF coil housing for each measurement. Planning of the MRSI slice position was aided by visual comparison of anatomical markers based on a sagittal 2D FLASH sequence which was acquired before each of the two ^1^H MRSI scans.

#### 2.1.3 Scan Protocol

Ten 1H MRSI slices were acquired in all acquisitions. The total FOV (220×220×80 mm^3^) was divided into three blocks of individual transversal slices (220×220×8 mm^3^): the top and middle blocks both contained four slices each, and the bottom block contained only two slices. Figure 1a shows the positioning of the three blocks. The uppermost block was placed immediately above the corpus callosum, and the two blocks below were positioned to extend the FOV based on the top block position.

**Figure 1:**
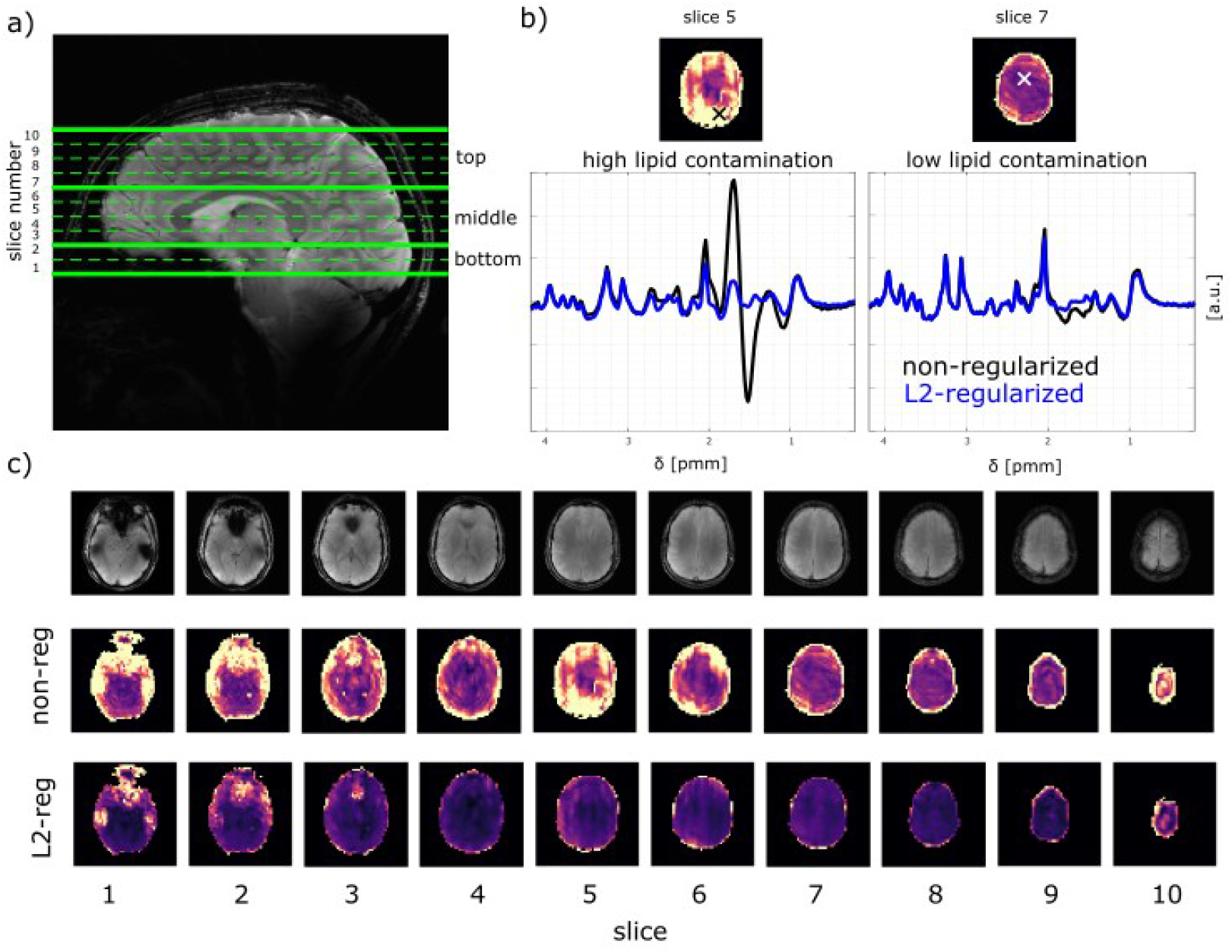
a) Anatomical image showing the slice positions of the 10 measured slices (total FOV 220 × 220 × 80 mm3) and the position of the three blocks of slices used for block wise B0 shimming. b) Sample spectra from a region with low and with high lipid contamination are shown from slice 5 and slice 7, respectively. The crosses indicate the origin of the sample spectra in the lipid maps of the non regularized data. Spectra without regularization (black line) and L2-regularization (black and blue line, respectively) are displayed. c) Anatomical images and respective lipid maps of each slice are shown for the data without regularization and with L2-regularization for volunteer V5.

Each block was B0 shimmed separately using the vendor implemented image-based second-order B_0_ shimming routine and a shim volume that extends an additional 4 mm beyond the bottom and top of each block of MRSI slices. The spectral data was circularly sampled using a 2D FID-MRSI sequence^31,33,46^ with a numerically optimized 3-pulse water-suppression scheme^29^, but without any lipid or outer volume suppression. MRSI slices were acquired with a matrix size of 48×48 (nominal voxel size: 4.6×4.6×8mm^3^=170*μ*L), TR=300ms, TE*=1.5ms, flip angle=48°, spectral width=8000Hz including oversampling, acquisition time=128 ms. After MRSI data acquisitions for each block, a non-water suppressed reference scan was acquired with the same parameters as the MRSI scan, but using a highly accelerated compressed sensing scheme with an acceleration factor of R~18^28,47,48^ to meet the scan time restrictions given by the local ethical committee. Additionally, a 2D FLASH scout image was acquired with TR=300ms, TE=6.1 ms, flip angle=25°, base resolution: 128×128 for the SENSE x-t sparse compressed sensing (CS) reconstruction of the water reference^28,47,48^. The scan time for the water suppressed MRSI data, the water reference MRSI data, and scout image accumulated to 10 min per slice. The total scan time summed up to 2h including slice positioning and shimming.

### 2.2 Data Processing and Postprocessing

An overview of the different reconstruction steps is shown in Table 1 and steps 1 to 7 are described in more detail in the supplementary material.

**Table 1:** Processing, post-processing and LCModel steps together with the quality metrics used for the different steps to optimize the procedure.

| Methods |  | Steps |  |  |
| --- | --- | --- | --- | --- |
| **Reconstruction and Processing** |  | 1. CS reconstruction of the accelerated water reference (wref)<br>2. Hanning filtering of the MRSI and wref data<br>3. spatial Fourier transformation of the MRSI and wref data<br>4. eddy current and phase correction of the MRSI and wref data using the wref data<br>5. coil combination using SVD of the MRSI and wref data |  |  |
| **Post-Processing** |  | 6. water removal using HSVD of the MRSI data<br>7. missing point prediction of the MRSI data<br>8. lipid regularization: |  |  |
|  |  | **No regularization** |  | **L2-regularization**<br>(same regularization parameter for all slices, derived from L-heuristic) |
| **LCModel Fitting** | **Macro-Molecular spectra** | Without macromolecule and lipid spectra | With simulated macromolecular spectrum | With simulated macromolecular spectrum |
|  | **LCModel stiffness parameter** | 0.15 (LCModel default)<br>0.25<br>1 |  |  |
| **Configuration Name** |  | nonreg-0.15<br>nonreg-0.25<br>nonreg-1 | nonreg-MM-0.15<br>nonreg-MM-0.25<br>nonreg-MM-1 | L2-MM-0.15<br>L2-MM-0.25<br>L2-MM-1 |
| **Quality Metrics** |  | • SNR of NAA and Cr, FWHM<br>• percentage of voxels with CRLB < 40 % for different metabolites<br>• root-mean-square of the fit residual<br>• Coefficient of variance for different metabolites<br>• 2D/3D maps |  |  |

The following sections explain the detail about the post-processing steps and the LCModel fitting procedure.

#### 2.2.1 Lipid suppression by L2-regularization

Following the post-processing, the data stayed either as it is (*“non-regularized”/“nonreg”*) before spectral fitting or a retrospective lipid suppression method via L2-regularizion (“*L2-regularized”/“L2”*) using an adaptation of Bilgic’s code^49^ (β=5000, obtained from L-curves^50^) was applied to reduce lipid contamination. Detailed information about the choice of the regularization parameter β can be found in the Supplementary Material and in Supplementary Figure A.

#### 2.2.2 Spectral fitting

The reconstructed MRSI data was fitted using LCModel (V6.3-1L)^51^ with a basis set simulated with VeSPA (version 0.9.5 https://scion.duhs.duke.edu/vespa/52) consisting of 13 metabolites: Asp, tCr, GABA, Gln, Glu, glycine (Gly), GSH, mI, NAA, NAAG, scyllo-inositol (scyllo), Tau, tCho (glycerophosphocholine + phosphorylcholine). J-coupling constants and chemical shifts were taken from Govindaraju et al.^53,54^ except for GABA for which Near’s values were used^55^. The LCModel parameter dkntmn (minimum spacing of the spline baseline knots in ppm, cannot exceed one third of the fitted range^56^) reflecting the spline baseline stiffness was set to 0.15 (default; most flexible), 0.25 (intermediate flexibility) or 1 (stiff) for comparison. The parameters of the LCModel control file and LCModel’s basis set are attached in the Supplementary Material. In addition, two sample spectra for a gray matter (GM) rich and a white matter (WM) rich voxel with all fitted metabolites are shown in the Supplementary Figure B.

The short TR times used in this work led to strong T1-weighting of metabolites and macromolecules (MM). To account for MM contributions, a simulated MM spectrum (MMAXIOM in Wright et al.^57^) was used as part of the spectral basis set. This model considered T1- and T2-relaxation times of MM at 9.4T^58,59^ and took into consideration the TR and TE of the 1H MRSI sequence. LCModel’s internal simulated MM or lipids were not included in the fitting. The relaxation corrected MM simulation model was used herein because it is not possible to acquire matching experimental macromolecular spectra with the same short TR and hence same T1 weighting as in the ^1^H MRSI measurements due to the need of a dual inversion recovery module^60^. Data fitted with the MM spectrum was compared to that without MM.

The following different configurations were compared: non-regularized data fitted without MM spectrum with dkntmn of 0.15, 0.25 and 1 (designated as *nonreg-0.15, nonreg-0.25* and *nonreg-1*, respectively), non-regularized data fitted with the simulated MM basis spectrum (designated as *nonreg-MM-0.15, nonreg-MM-0.25* and *nonreg-MM-1*, respectively), and L2-regularized data fitted with the MM basis spectrum (designated as *L2-MM-0.15, L2-MM-0.25* and *L2-MM-1*, respectively).

### 2.3 Quality Measures & Test-Retest reliability

Different quality measures were used to compare the mentioned configurations and choose the best one as input for the subsequent test-retest analysis.

The contribution of skull lipid signal to the metabolite spectrum can elevate apparent metabolite peak amplitudes while a wrong adjustment of the regularization parameter in the retrospective lipid removal step can also suppress metabolite signal. To quantify this impact, the signal-to-noise ratio (SNR) of NAA and tCr were observed. Thus, the SNR of NAA and tCr were calculated as the ratio of the maximum peak of their methyl groups at 2 or 3 ppm derived from the LCModel fit to the root-mean-square (rms) of the noise region of the non-back-predicted ^1^H FID MRSI data (to avoid baseline distortions caused by the back-prediction). In addition, a 2nd order polynomial fit of the baseline was subtracted prior to the noise level estimation. Using the maximum of the fitted metabolite spectra as the signal intensity allowed to account for baseline compensations from LCModel and the choice of different LCModel baseline stiffness’.

Furthermore, the full width half maximum (FWHM) linewidth of tCr as reported from LCModel, the root-mean-square (rms) of the fit residual and the Cramér-Rao lower bound (CRLB) as reported from LCModel were reported for all configurations. In addition, the brain coverage of different metabolites by using the percentage of voxels fitted with a CRLB < 40 % to compare it to previous published data^38^, is shown.

To investigate the intra-subject reliability of the two measurements the coefficient of variance (CV), defined as standard deviation of the difference between measurements divided by the mean values from the measurements^11^, were calculated for NAA/tCr, tCho/tCr, Glu/tCr, and mI/tCr with the mean concentration of each slice and for NAAG/tCr, GABA/tCr, GSH/tCr, Tau/tCr, Gln/tCr, Asp/tCr for slice 6-9 due to the insufficient quality in lower slices.

Two-sample *t*-tests with non-equal variances were used to prove statistical significance at the 5% significance level of the above mentioned quality measures for different configurations. Whenever changes are described as significant, this two-sample t-test was applied.

## 3. Results

### 3.1. Lipid Suppression

The lipid rms maps of non-regularized data show strong lipid contamination especially in slices 5 and 6, see Figure 1c. The impact of the regularizations on the spectra is shown in Figure 1b for strong and intermediate contaminated regions.

### 3.2 Signal-to-Noise-Ratio

Boxplots of the SNR of NAA and tCr for each configuration are displayed in Figure 2a+b. This reveals that the regularization influences different spectral ranges with varying effects: While the SNR of NAA decreases slightly after the regularization, the SNR of tCr changes less. These changes are statistically significant for all slices combined as well as for the single slices. A nonregularized configuration and a L2-regularized configuration are selected to illustrate the changes in different slices (Figure 2c+d): The SNR increases for slices 1 to slice 5, show decreased values for slice 6 and show highest values for slice 7 and 8.

**Figure 2:**
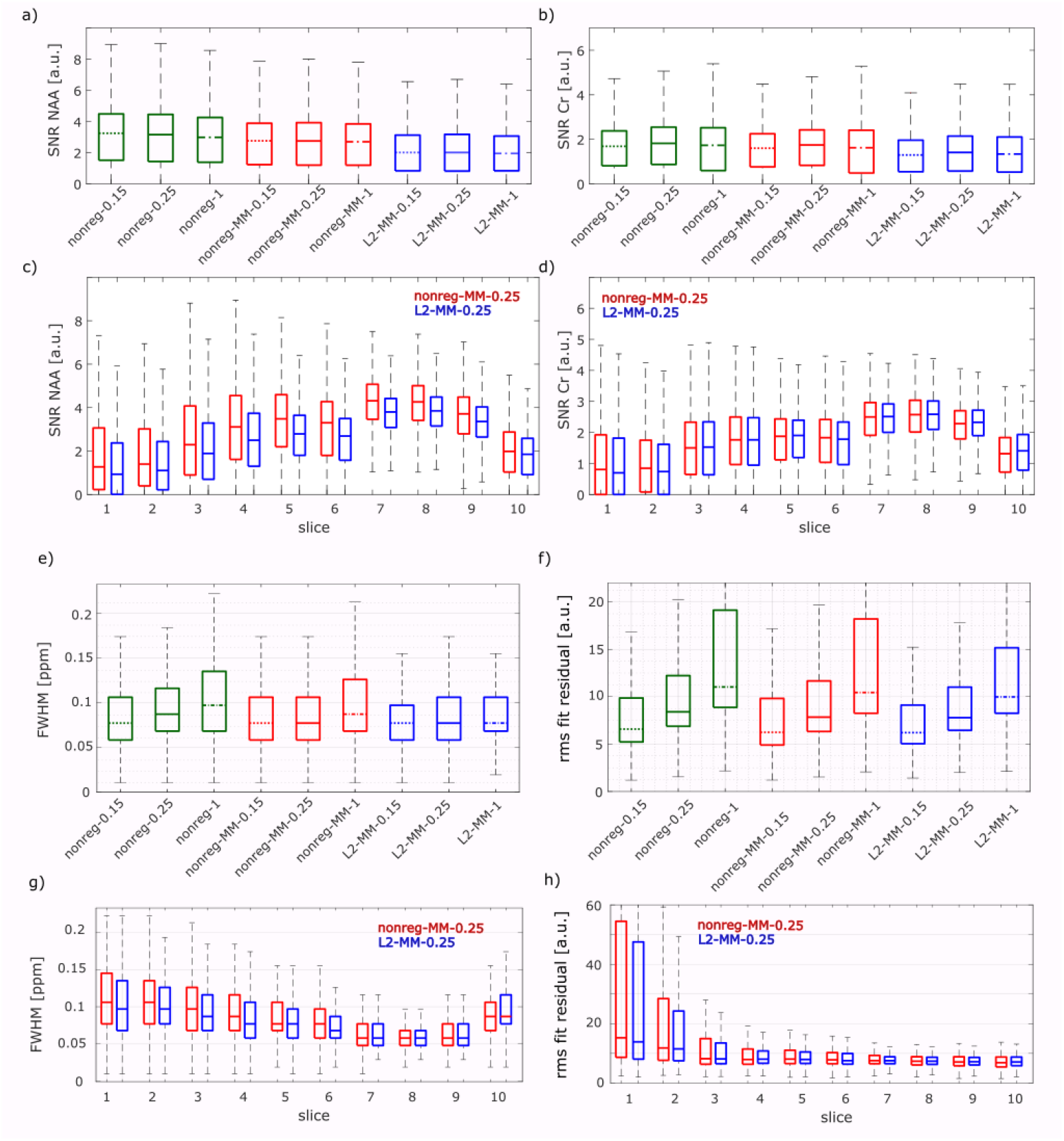
Boxplots of the SNR of NAA (a) and Cr (b), FWHM of Cr (e) and the rms fit residual (f) are shown for all configurations without and with L2-regularization (nonreg vs L2), with and without simulated MM spectrum and with different LCModel baseline stiffness values (0.15, 0.25 and 1). Data from all measurements and volunteers are pooled. Corresponding slice specific values pooled across measurements and volunteers from the configurations “nonreg-MM-0.25” (red) and “L2-MM-0.25” (blue) are seen in c,d,g,h). In all boxplots, the horizontal line indicates the median and the boxes show the [25-, 75]-percentile interval.

### 3.3 Fitting Parameters

The FWHM as well as the rms of the fit residual increase with increasing dkntmn as it is seen in Figure 2e and 2f, respectively. The FWHM changes significantly. The rms fit residual are slightly smaller for all L2-regularized cases than for the non-regularized configurations. These changes are statistically significant, while the rms fit residual for non-regularized data does not change significantly for the data with and without MM spectrum for the same dkntmn value. The fit residual can be seen in the Supplementary Figure C for one sample voxel for all configurations.

The CRLB increases with increasing dkntmn value for all shown metabolites, see Figure 3a and Supplementary Figure D. The same behavior is seen in the voxels fitted in CRLB < 40 %, see Figure 3b.

**Figure 3:**
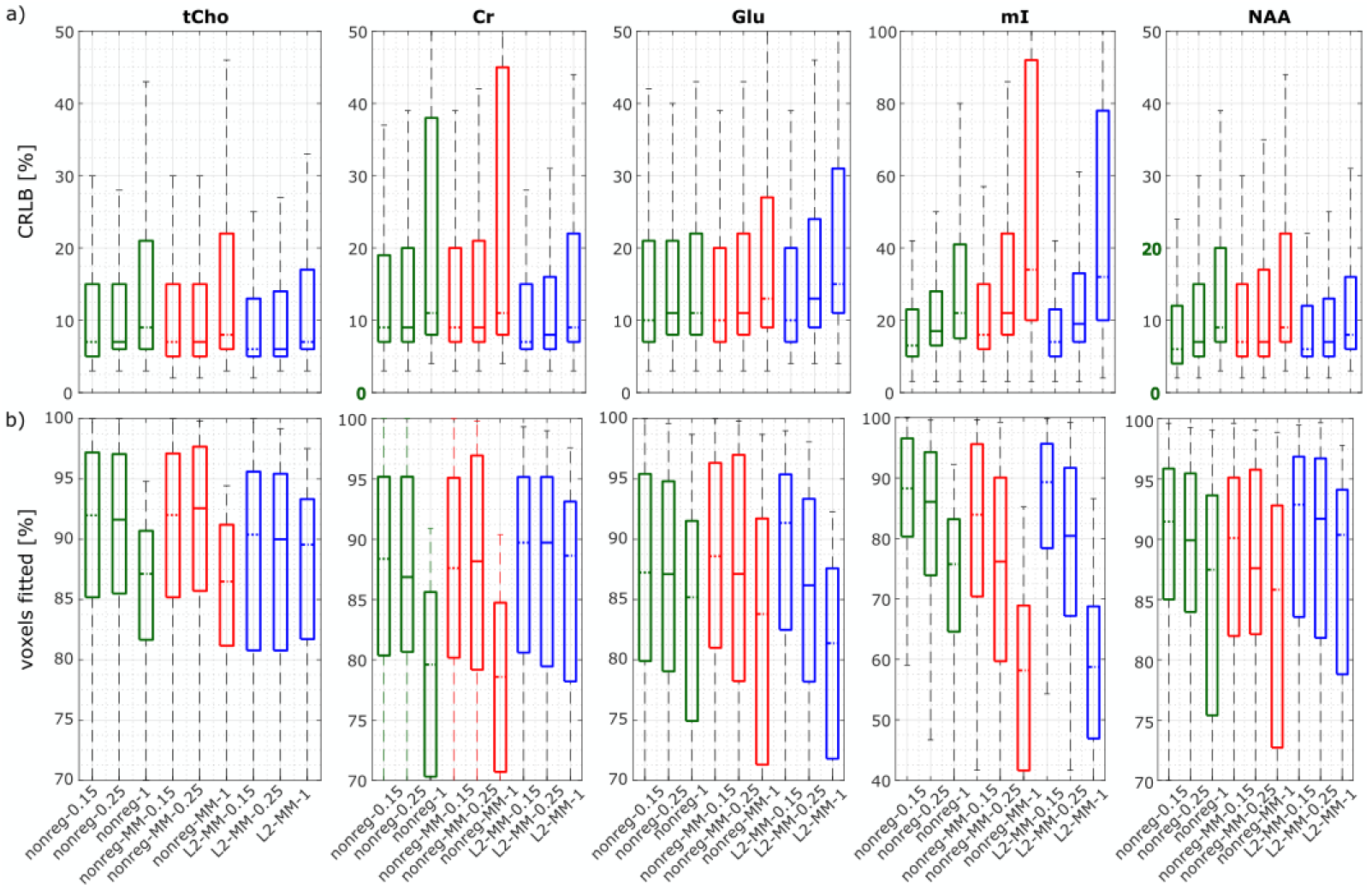
CRLB (a) and voxels fitted (b) from LCModel for all configurations and different metabolites. Data from all measurements, volunteers and slices are pooled. CRLB and percentage of voxels fitted for the other metabolites can be found in the Supplementary Material Fig D.

### 3.4 Qualitative Analysis

In Figure 4, metabolite ratio maps are shown for selected slices for Glu/tCr (a), NAA/tCr (b), NAAG/tCr (c) for all configurations to illustrate the effects of different dkntmn values, MM spectra and lipid regularization. tCr ratios were used to include B1^+^ and B1^-^ correction^31^. In Figure 5, metabolite ratio maps for the sagittal and coronal slices (a) for NAA/tCr, tCho/tCr, Glu/tCr and mI/tCr is shown as well as all measured transversal slices (b) for one volunteer for 10 different metabolites of the configuration *L2-MM-0.25*.

**Figure 4:**
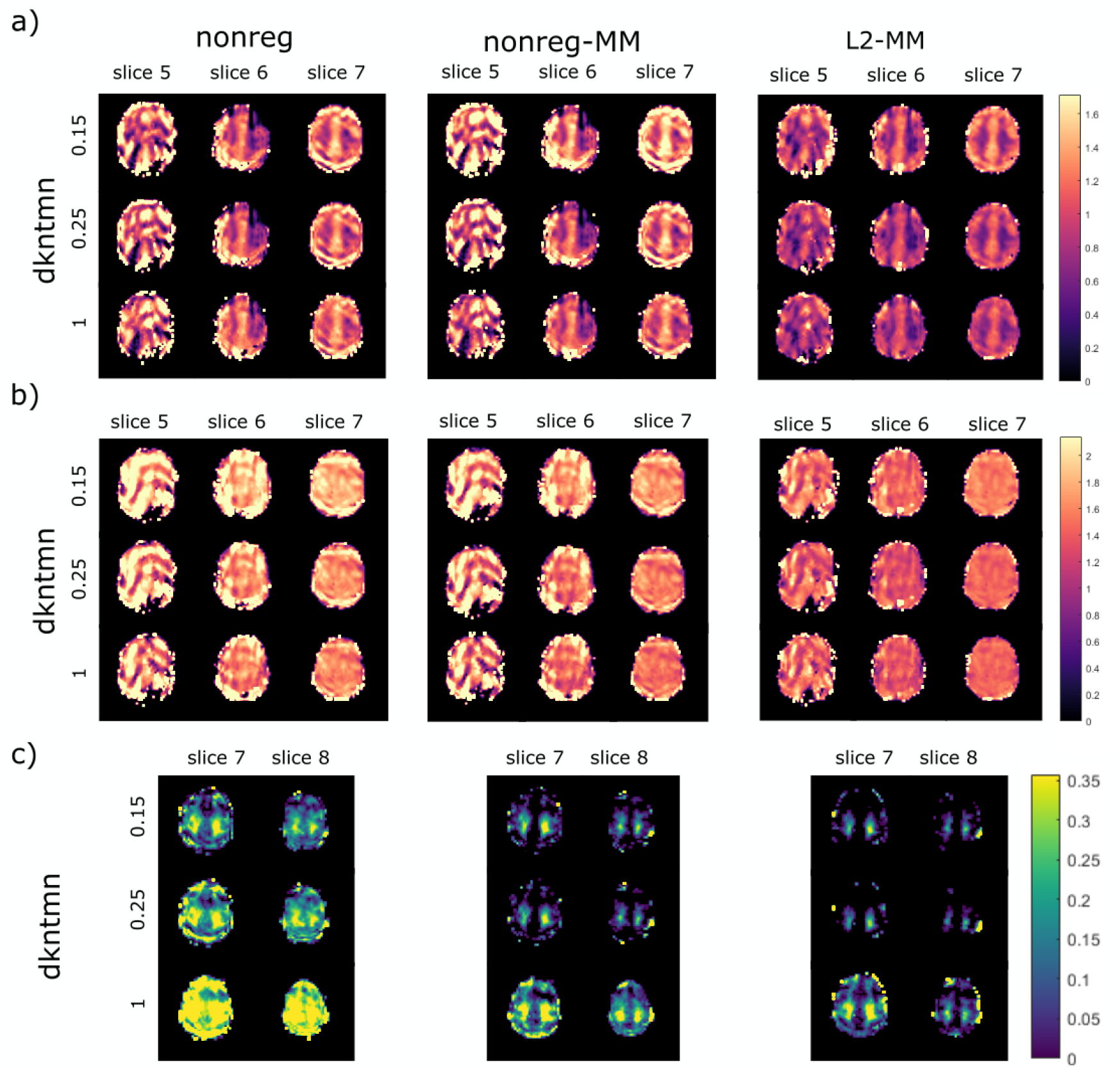
Normalized concentration maps in abitrary units for Glu/tCr (a), NAA/tCr (b), NAAG/tCr (c) for selected slices from volunteer V5 for all configurations. Be aware of the different color scale for NAAG due to better visibility of small concentrations. Both color scales are linear.

**Figure 5:**
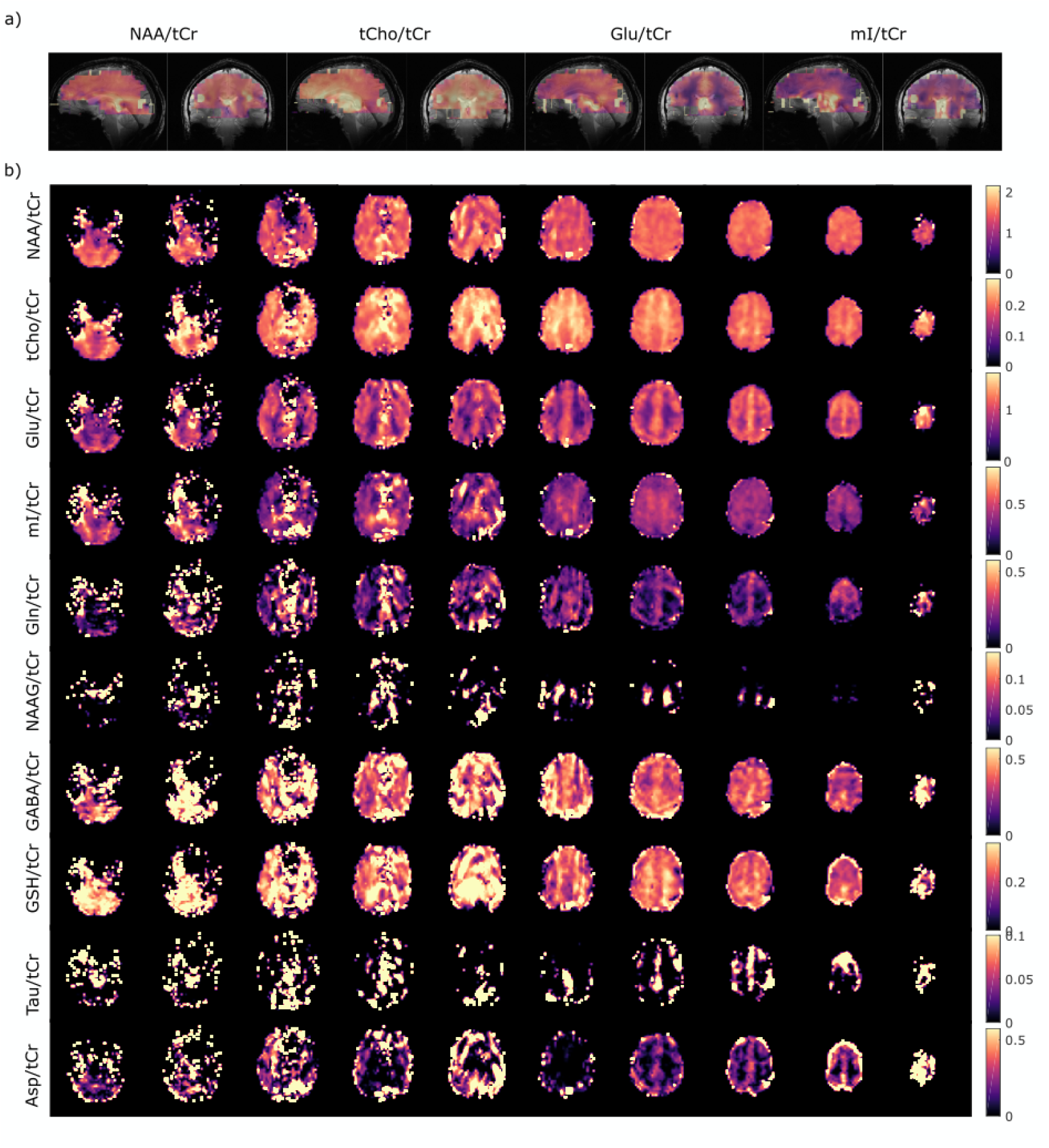
a) Sagittal and coronal metabolite ratio maps (/tCr) for four selected metabolites, underlayed with the anatomical images to illustrate the coverage of the cerebrum. b) Transversal metabolite ratio maps for all slices. Data from configuration L2-MM-0.25 and volunteer 5. Data reported in arbitrary units.

### 3.5 Test-retest Analysis

In Figure 6, test-retest ratio maps are shown for all 10 metabolites of the configuration *L2-MM-0.25*. In Table 2, the CV for the same metabolites were summarized, the corresponding boxplots including 25-und 75 % percentile can be found in the Supplementary Figure E.

**Figure 6:**
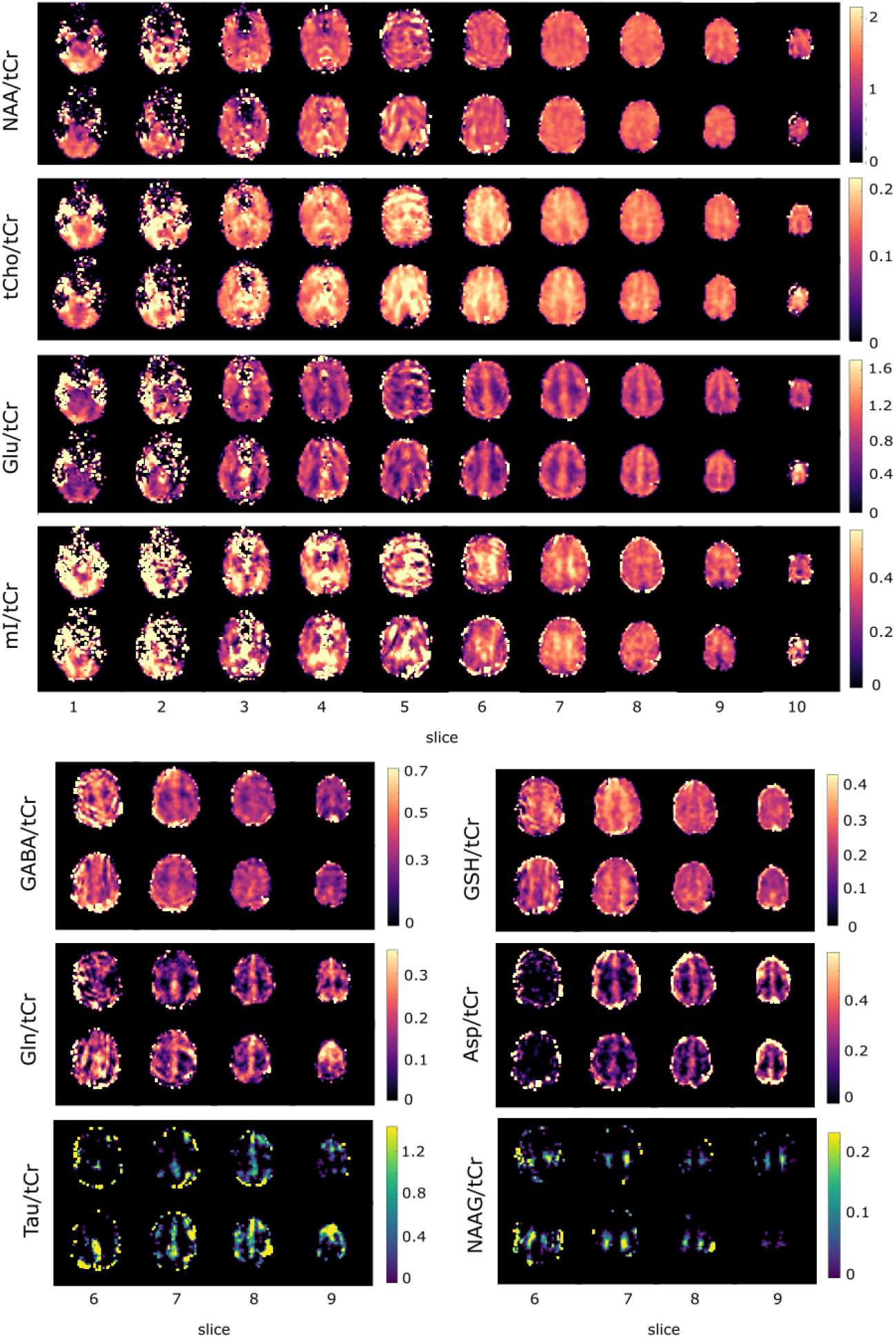
Metabolite ratio maps for volunteer V5 for both measurements to present the reproducibility of the measurement; configuration L2-MM-0.25 data. Data reported in arbitrary units. For four metabolites, only a subset of slices is shown due to unreliable results in lower slices.

**Table 2:** Slice-wise intra-subject coefficient of variations (CVs, mean, std in %) for ten normalized concentration ratios (/tCr) for six configurations for all volunteers and either all slices (#) or slice 6-9 (*).

| Metabolite (/tCr)<br>vs. Method | NAA<sup>#</sup> |  | tCho<sup>#</sup> |  | Glu<sup>#</sup> |  | ml<sup>#</sup> |  | Asp<sup>*</sup> |  |
| --- | --- | --- | --- | --- | --- | --- | --- | --- | --- | --- |
|  | mean | std | mean | std | mean | std | mean | std | mean | std |
| nonreg-0.15 | 5.6 | 3.3 | 5 | 3.2 | 7.4 | 4.2 | 7.2 | 2.9 | 14.0 | 8.6 |
| nonreg-0.25 | 6.3 | 3.6 | 5.1 | 3.2 | 8.4 | 4.6 | 9.0 | 3.4 | 16.0 | 7.7 |
| nonreg-1 | 10.8 | 5.9 | 12.6 | 16.4 | 14.3 | 8.9 | 20.0 | 8.2 | 23.3 | 11.5 |
| nonreg-MM-0.15 | 6.2 | 3.3 | 4.6 | 2.7 | 6.5 | 4.2 | 7.4 | 2.9 | 15.4 | 5.0 |
| nonreg-MM-0.25 | 6.3 | 3.6 | 5.1 | 3.4 | 8.3 | 5.2 | 12.8 | 1.1 | 15.8 | 7.0 |
| nonreg-MM-1 | 9.1 | 5.4 | 12 | 14.1 | 14.4 | 10.8 | 29.1 | 13.4 | 25.3 | 18.0 |
| L2-MM-0.15 | 7.2 | 4.9 | 4.3 | 2.3 | 11 | 7.1 | 12.6 | 8.7 | 15.0 | 0.8 |
| L2-MM-0.25 | 8.5 | 4.7 | 4.1 | 2.2 | 10.3 | 6.2 | 16.7 | 8.8 | 17.3 | 5.5 |
| L2-MM-1 | 8.9 | 4.5 | 5.4 | 2.5 | 14.1 | 9.2 | 23.3 | 8.0 | 26.3 | 9.2 |
| Metabolite (/tCr)<br>vs. Method | GABA<sup>*</sup> |  | Gln<sup>*</sup> |  | NAAG<sup>*</sup> |  | Tau<sup>*</sup> |  | GSH<sup>*</sup> |  |
|  | mean | std | mean | std | mean | std | mean | std | mean | std |
| nonreg-0.15 | 8.0 | 3.6 | 14.3 | 8.7 | 15.6 | 4.2 | 21.1 | 11.3 | 4.9 | 2.0 |
| nonreg-0.25 | 9.2 | 4.3 | 15.9 | 9.0 | 16.5 | 3.6 | 24.4 | 10.3 | 4.2 | 2.4 |
| nonreg-1 | 11.1 | 3.3 | 27.9 | 28.2 | 14.4 | 5.4 | 51.2 | 38.1 | 9.9 | 5.4 |
| nonreg-MM-0.15 | 8.2 | 5.1 | 16.1 | 9.1 | 19.5 | 4.5 | 18.8 | 11.1 | 9.5 | 2.3 |
| nonreg-MM-0.25 | 9.0 | 4.6 | 22.1 | 13.9 | 20.8 | 7.2 | 22.0 | 8.6 | 10.5 | 3.7 |
| nonreg-MM-1 | 12.5 | 4.2 | 23.5 | 27.5 | 11.3 | 7.7 | 30.8 | 15.7 | 22.6 | 5.1 |
| L2-MM-0.15 | 10.6 | 6.4 | 17.7 | 13.1 | 19.5 | 12.5 | 19.2 | 20.2 | 6.2 | 2.5 |
| L2-MM-0.25 | 13.8 | 6.8 | 22.3 | 16.9 | 33.7 | 23 | 35.3 | 20.6 | 7.1 | 2.4 |
| L2-MM-1 | 14.5 | 5.1 | 28.0 | 19.3 | 12.2 | 5.2 | 89.4 | 28.9 | 12.6 | 2.2 |

## 4. Discussion

In the present study, non-accelerated multi-slice FID-MRSI data encompassing the cerebrum of five volunteers were acquired and analyzed with regard to the effects of an included MM spectrum and different values of the LCModel stiffness parameter in the fitting routine on nonregularized and lipid-regularized data. In addition, a second measurement on the same volunteer were performed and the effects of the mentioned configurations on intra-subject reliability were investigated.

In the next subsections, the calculated quality measures will be discussed with respect to the MM spectrum used in the fit, the influence of the retrospective lipid suppression and the choice of LCModel’s baseline flexibility. Finally, the metabolic ratio maps and the test-retest results will be discussed.

### 4.1 Influence of the MM spectrum

The correct estimation of the macromolecular signal is crucial for the accurate determination of metabolite concentration^60^. Several ultra-high field studies presented a measured full MM spectrum or decomposed MM signal into individual components for SVS^44,56,61^ as well as for MRSI data^42,62,63^. A simulated model is favorable in cases of limited scan time and when experimental acquired MM spectra cannot be obtained as it is the case in the present study. The present MM-model was successfully tested before on SVS data on 9.4 T^57^ for STEAM and semi-LASER, but is adaptable to different sequences, so also to FID-MRSI. In addition, it could also be adapted for different tissue types, but the impact was found to be minor^64^ and the experts’ consensus was that a constant shape of the MM can be a practical approach^60^. To observe the influence of the MM model on the present data, the non-regularized data fitted with and without a simulated MM spectrum (*nonreg* vs *nonreg-MM*) will be compared. The SNR of NAA and tCr is higher for *nonreg* configurations without the MM spectrum than with the MM spectrum for all dkntmn values, since MM peaks overlap with the peaks of NAA and tCr at 2 and 3 ppm, respectively. Thus, the fitted concentrations are reduced when using a MM baseline, which does not imply that the data quality is worse. The other quality measures (FWHM, the rms fit residual and the CRLB for tCho, NAA, tCr and Glu) did not differ significantly. Only the CRLB for mI was significantly higher when fitted with the MM spectrum which is caused since the MM spectrum and mI have both peaks around 3.5 ppm. The peaks are assigned completely to mI if no MM basis spectrum is used and to both if a MM basis spectrum is included which decreases the mI concentration and elevated the CRLB for mI. The ratio maps of mI/tCr do not show a quantitative improvement when including or not including the MM spectrum, see Supplementary Figure F.

Since these measures do not answer the question if the MM spectrum should be used or not, metabolite ratio maps were investigated qualitatively. A dominant impact of the MM spectrum was visually observed for NAAG/tCr, see Figure 4c. NAAG is expected to be much higher in the WM^33,63,65^ and the maps obtained in this study for dkntmn < 1 look much more similar to what has been published before^64^ when including the MM spectrum in the fit than without.

Consequently, it could be shown that the simulated MM-model could successfully be applied to FID-MRSI data, but in a final decision step, its impact on quantitative metabolic concentrations should be investigated, which is beyond the scope of this study.

### 4.2 Retrospective Lipid Suppression

Although, lipid suppression by L2 is widely used, there are only few publications beside the original article by Bilgic et al.^49^ (one paper combining water and lipid removal by Lin et al.^66^ and an abstract from Hangel et al.^67^) investigating the lipid removal by L2-regularization in more detail. All other MRSI publications at ultra-high fields using L2 (or rarely L1) give no hints on how they found the optimal regularization parameter and rarely report the value at all. Thus, it is essential to analyze the impact of L2 in more detail by comparing non-regularized data to L2-regularized data (*nonreg-MM* vs *L2-MM*).

On one hand, the lipid contamination can be efficacious reduced when using L2-regularization, see third row of Figure 1; however, on the other hand, it also strongly reduces the NAA CH_3_ peak height as seen in the sample spectra of Figure 1b. This can be partly traced back to a baseline distortion caused by the lipid regularization and partly to a peak reduction of peaks close to the chemical shift of the lipids (0.9 – 1.3 ppm), which results in stronger reduction of the SNR of NAA than the SNR of Cr, although both changes were statistically significant. The quality measures of the fitted data show no clear trend: The CRLB is partly significantly lower or higher and partly non-significantly different depending on the metabolite and the dkntmn value chosen; but the FWHM and the fit residual is significantly lower for all lipid-regularized cases and the number of voxels fitted with CRLB < 40 % is higher for most metabolites. Overall, the range of the median values for voxels fitted with CRLB < 40 % was calculated for the whole volume for tCr: 88-90 %, tCho: 90-93 %, Glu: 86-91 %, mI: 76-89 % and NAA: 88-98 % for the configurations *nonreg-MM* and *L2-MM* with dkntmn = 0.15 and 0.25. Higher dkntmn values lead to lower % of voxels fitted and were not reported here. These values are similar to the values reported from Hingerl et al^38^ in a 3D-MRSI study from the highest quality region: tCr: 91 %, tCho: 93%, Glu: 91%, mI: 92 %, and tNAA: 94%.

To decide finally whether to use the lipid suppression or not, the NAA/tCr and Glu/tCr maps will be inspected, since they show the strongest effects of lipid suppression of all investigated metabolites, see Figure 4a+b. It turns out that lipid suppressed data reduces the artifacts clearly, which are present in the non-regularized data (no matter which dkntmn values was used). Especially the artifacts in slice 5 were reduced (for both metabolites), but also the data loss in slice 6 in the Glu/tCr maps could be removed and the increase of the NAA/tCr concentration in peripheral voxels in slice 6 and peripheral artifacts in slice 7 for both metabolites cannot be detected in the L2-regularized data. Hence, while L2-regulization reduces artifacts effectively, it does not create new artifacts as it can be seen for all other slices for NAA/tCr and Glu/tCr in Supplementary Figure G.

### 4.3 Spline Baseline Flexibility

No previous literature systematically analyzed the influence of different LCModel’s dkntmn values on metabolite maps. Most papers do even not report the used dkntmn value. There is one study on baseline problems using a different fitting tool for SVS and MRSI data^43^ showing that baseline-fitting errors can be drawn back to too stiff and too flexible baselines leading to underfitting and overfitting, respectively. Since LCModel is most widely used, it is crucial to systematically investigate its internal spline baseline flexibility.

There are several SVS studies which are comparing different dkntmn values, but those results should only be adapted with caution since the baseline distortion is stronger for MRSI than for SVS data. Previous studies on human SVS data show that LCModel’s default baseline stiffness (0.15) may be to flexible and encourage stiffer baselines^56,61,68,69^ to avoid quantification errors due to fitting ambiguity, but no conclusion on a best values was drawn. A SVS study in rat brain from Simicic et al^70^ concluded that a dkntmn value < 1 is preferred when comparing the concentrations to a ground truth. But no generally valid value for dkntmn was predicted since it is i.a. depending on the MM spectrum used.

For the MRSI data in the present study, it can also be concluded that a very stiff baseline (dkntmn=1) is not a good choice: The SNR values are lowest, the FWHM highest, the rms of the fit residual increased and the CRLB are highest and the number of voxels fitted thus lowest. The NAAG maps show obviously anatomically incorrect results. Although the LCModel’s default baseline stiffness (dkntmn=0.15) show in most cases better values of the quality metrics’ than a dkntmn value of 0.25, it has to be kept in mind that an overflexible baseline potentially leads to quantification errors^61^. Further investigations including the comparison of quantitative metabolite concentrations for different dkntmn values should be carried out to find an optimal baseline stiffness. But a dkntmn value close 0.25 is potentially a preferred choice for MRSI data. In MRSI papers values of 0.15^42^ and 0.3^71^ were found.

### 4.4 Metabolite Ratio Maps

In the previous subsection, it has been shown that L2-regularized data fitted with a flexible baseline and a MM prior knowledge is the best choice. Thus, the configuration *L2-MM-0.25* was chosen for presenting metabolite ratio maps in Figure 5. Since no tissue segmentation was done in this study, the concentration ratio maps can only visually be compared to previous literature, but correspond mostly well to the expected ratios of GM and WM: higher concentration in WM than GM for tCho/tCr^21,37–40,46^, mI/tCr^39,46^, NAAG/tCr^31,39,63,65^ and higher concentrations in GM for Glu/tCr^31,37,38,40,46^, GABA/tCr^31,72,73^, Gln/tCr^31,74^, Tau/tCr^31,74^, Asp/tCr^31^. For some metabolites even more anatomical details are seen in the maps which correspond to the literature: NAA/tCr is higher in the corpus callosum and thalamus^37^; tCho/tCr is concentrated higher in the frontal cortex than in the occipital lobe^21,38–40^ and strongly in corpus callosum and thalamus^37^. In the literature NAA/tCr is higher in WM than GM^21,31,39,46,63^, while the transversal concentration ratio maps in Figure 5b show a rather evenly distributed NAA/tCr concentration and look similar to the maps presented in other publications^12,31^. Also the values for GSH/tCr were difficult to compare: Previous literature show higher GSH concentration in GM^75^, but for the concentration ratio, literature results are not consistent: GSH/tCr is found to be similar in GM and WM^71^, higher or lower in GM depending on the resolution^31^ or slightly higher in WM than in GM^37^, but our values are definitely higher in WM. A possible reason could large signals from other metabolites overlap with GSH which hampers its fitting^76,77^.

Overall: the best results were achieved in the upper slices (slice 6-9, corresponds to a slab of 32 mm thickness) for all volunteers, which can be easily understood since the shim is best in this region: the FWHM is smallest, see Figure 2g, and SNR is biggest, see Figure 2c+d. In this volume tCho, NAA, Glu, mI, Gln, GABA, GSH, Tau, Asp and NAAG show good quality metabolite ratio (/tCr) maps. Slice 10 covered only a very limited number of brain voxels and thus, the data of the top slice was strongly contaminated by the surrounding lipid from the skull. No information could be obtained for this slice for any volunteer. The slices of the lowest block (slice 1 and 2) suffer from static magnetic field (B_0_) inhomogeneity caused by the nasal sinus in the frontal region. Depending on the volunteer’s head size, shape and relative orientation to the outer magnetic field information of only a few metabolites (NAA, tCr, tCho, Glu and mI) could be achieved in these lower brain slices. Slices 3-5, too, suffer from the mentioned B_0_ inhomogeneity and reasonable concentration ratio maps could be achieved for some volunteers for NAA, tCr, tCho, Glu and mI.

Overall, in some metabolite ratio maps other artifacts especially in slice 5 are still present after L2-regularization, which could either be caused by the higher peripheral and lower central receive sensitivity of overlapping coil elements (see Figure 1 in Avdievitch et al.^45^). Other artifacts could be explained by motion during the long duration of the scan. Motion would have diminished the B0 homogeneity achieved by the B0 shimming procedure and thus reduced the quality of the MRSI data. In addition, motion in between the MRSI data and the water reference acquisition (time difference of 10-40 minutes), could have created mismatches between the slice position of the water suppressed and non-water suppressed data.

### 4.5 Inter-session, Intra-subject reliability

Although there are numerous test-retest studies of MRSI^13,22,25,26,42,78,79^ (and even more for SVS^80–90^) data, comparing their CVs to the present data is challenging since many do not show CVs of concentration ratios but of the absolute concentration and secondly, often voxelwise CVs are reported averaged from different brain regions or tissue types, while in the present study CVs were calculated slicewise. Nonetheless, the most comparable studies will be cited in the next paragraph beginning with (towards) whole-brain studies. Single-slice MRSI as well as SVS reproducibility CVs were only cited as a rough guide for the CVs of those metabolites, which are not reported in whole-brain studies.

In the present study the mean inter-session, intra-subject CV for the configurations *nonreg-MM* and *L2-MM* using dkntmn values of 0.15 or 0.25 ranges from 6.2-8.5 %, 4.6-5.1 %, 6.5-10.3 % and 7.4-16.7 %, for NAA/tCr, tCho/tCr, Glu/tCr and mI/tCr for the whole volume, respectively. And the CVs for slice 6-9 were found to be 15.0-17.3 %, 8.2-13.8 %, 16.1-22.3 %, 19.5-33.7 %, 18.8-35.3 %, 6.2-10.5 % for Asp/tCr, GABA/tCr, Gln/tCr, NAAG/tCr, Tau/tCr, GSH/tCr, respectively. To our best knowledge, there are no reproducibility studies at ultra-high fields encompassing a larger volume in the brain. There are a few other studies reporting inter-session, intra-subject CVs at lower fields for concentration ratios (/tCr) of NAA, tCho, mI and Glx. Their values are similar to those reported here: Zhang et al.^26^ calculated mean intra-subject CVs from 3 sessions in different regions from whole-brain EPSI data at 3T of NAA/tCr: 6%, tCho/tCr: 6.5 % and mI/tCr: 10.5 %. Maudsley et al.^22^ calculated mean intra-subject CVs from 5 session acquired with SE MRSI at 3T ranging from NAA/tCr: 1.8-11 % and tCho/tCr: 2.7-10.2 % for different regions and Gu et al.^13^ reported from 6 scans acquired with 3D MRSI ranging from NAA/tCr: 2.3-12.7 % and tCho/tCr: 2.0-7.1 % for different regions at 1.5 T. Hnilicova et al.^79^ calculated CVs of Glx/tCr of 4-8% and GABA+macromolecules/tCr of 6-10% (minimal-maximal value of medians) for using MEGA-LASER 3D MRSI sequence at 3T. Others reported intra-session or inter-subject CV and will not be compared^11,23,25,37^. Due to the lack of comparable high-field reproducibility studies with large volumes with more metabolites, the CVs of a single-slice study at 7T from Heckova et al.^42^ will be referenced. They showed voxel-by-voxel single-slice CVs for different brain regions of tNAA/tCr: 7 %, tCho/tCr: 5.9 %, mI/tCr: 7 %, Glx/tCr: 8.1 %, Glu/tCr: <12%, GSH/tCr: <15%, and Gln/tCr and Tau/tCr of <20%. These values are in a similar range as the present data with the CVs of mI/tCr and Tau/tCr being slightly smaller, and GSH/tCr being slightly higher than in our results. CVs for Asp and NAAG could not be found in other MRSI studies. In SVS studies values of mean CVs for NAAG/tCr for the posterior cingulate cortex of 24.8%^83^ at 7T, which is in the middle of our CV values for different configurations. Reference CVs of Asp/tCr could not be found.

Except for tCho, the CV values show better results for nonregularized data than for L2-regularized data. It is thus concluded that as long as no lipid artifacts are visually seen, nonregularized data might be the better choice.

### 4.6. Limitations and future work

A recent review paper summarizes all previous publications about accelerated MRSI (> 200) and includes recommendations on which of them are applicable to 1H MRSI at ultra-high field^91^. A commonality between all previous ultra-high field “whole-brain” ^1^H MRSI studies^36–40^ is that they used acceleration methods to achieve (towards) extended brain coverage in short scan times. In contrast, the aim of the current study was to use a non-accelerated approach to achieve high spectral and spatial resolution metabolite maps for 10 brain metabolites that shall serve as a reference for neuroscientific and clinical studies and also as a ground truth for future analysis on the impact of different acceleration methods.

In future studies, the B0 shimming performance should be further improved to obtain better quality spectra in lower brain regions and to reduce remaining artifacts especially in the middle area of the measured volume (around slice 5). However, in spite of the investigation of different hardware and software solutions this is a problem so far not solved by the entire MRI developer community^1,92^. In addition, the reduction of motion artifacts is needed by prospective motion correction as well to make whole brain 1H MRSI at ultra-high field more appropriate for clinical studies.

## 5. Conclusion

In this study, reproducible concentration ratio maps (/tCr) of tCho, NAA, Glu, and mI for the whole cerebrum and of NAAG, GABA, GSH, Tau, Gln, Asp for a volume of 32 mm thickness were obtained for five volunteers with good correspondence to the anatomical structure using fully sampled multi-slice 1H FID MRSI data at 9.4T with retrospective L2-lipid-regularization, an optimized flexible fitting baseline and a simulated-MM model.

## List of Supporting Information

Supporting Information to the Method Section (Text)

Figure A: L-curve (lipid basis norm vs. L2 of consistency) and the corresponding curvature (b) from one sample slice. Curvatures from different slices and volunteers (c).

LCModel.control File:

Figure B: Sample spectra from dominant gray matter (a) and white matter (b) voxel with all fitted metabolites for the configuration L2-MM-0.25.

Figure C: Data, fit, MM spectrum, residual and spline baseline for each configuration from one sample spectrum of volunteer V5.

Figure D: CRLB (a) and voxels fitted (b) from LCModel for all configurations and different metabolites. Data from all measurements, volunteers and slices are pooled.

Figure E: Coefficient of variance of the metabolite ratios (/tCr) for the test-retest measurements for all configurations and ten metabolite concentration ratios. Data pooled from all volunteers and a) all slices, b) slice 6-9.

Figure F: mI/tCr concentration ratio maps for the configurations: nonreg-0.15, nonreg-0.25, nonreg-1 and nonreg-MM-0.15, nonreg-MM-0.25, nonreg-MM-1 for volunteer 5. Data reported in arbitrary units.

Figure G: Comparison of metabolite ratio maps (NAA/tCr and Glu/tCr) for different regularization parameters: First row shows non-regularized data, the rows below show L2-regularized data for β=100, 1000, 5000, 10000 and 106, respectively.

LCModel Basis Set

