## Supplementary for "Test-retest reproducibility of human brain multi-slice 1H FID-MRSI data at 9.4 T after optimization of lipid regularization, macromolecular model and spline baseline stiffness"

### Supporting Information to the Method Section

In the following supporting information to the method section, step 1 to 7 of the processing and post-processing steps listed in table 1 is described in more detail.

Water Reference MRSI Data Reconstruction (Step 1)

Missing k-space points of the highly undersampled (R~18) water reference MRSI data were calculated using the CS algorithm of Nassipour et al.[1–3], which included SENSE reconstruction based coil sensitivity ESPIRiT maps derived from the 2D FLASH scout image prior to the CS reconstruction step.

Reconstruction and Processing Methods (Step 2-5)

Metabolite and water MRSI data were processed using the following methods: Hanning filtering, 2D spatial Fourier transform, eddy current and $0^{th}$ order phase correction using the water reference data and coil combination by singular value decomposition (SVD)[4]. Additionally, the dc-offset was removed as well as the residual water signal using the Hankel-Lanczos method (HLSVD)[5].

Post-processing Methods (Step 6 and 7)

Due to the acquisition delay of 1.5ms, a first-order phase was induced in the data. This phase was corrected by missing-point prediction using linear back prediction with the auto-regressive Burg method[6]. No further apodization, zero-filling or additional phase correction was applied.

Lipid-regularization (Step 8)

L2-regularization was used to reduce artifacts from lipid contamination, which occurred in some slices. An adaption from Bilgic's code[7] was used with lipid maps defined as all voxels with the lipid root mean square (rms) of >8 [a.u.]. This factor was chosen by visual inspection to get lipid ring maps reflecting the high lipid content of the skull and skin for all volunteers and slices. Brain masks were calculated as all voxels within the respective lipid ring. Due to the lack of additional data with more averages and different spatial resolutions – as it was needed for the original code from Bilgic – no dual density image was calculated before the regularization. The regularization parameter β war calculated for each slice and volunteer separately using the L-curve criterion[8]. In Figure A, a sample L-curve (a) and the corresponding curvature (b) is shown for one sample slice. In Figure Ac) several curvature curves are displayed. Unfortunately, the calculation of the optimal β was not successful for each slice. So, the effects of different β in metabolite maps were compared to obtain the optimal value for our data. In Figure G, metabolite ratio maps for NAA/tCr and Glu/tCr are presented for different β. Evaluating these maps, the regularization parameter was set to an intermediate value of those slices which could be evaluated (β=5000), which seems to reduce lipid artifacts, but is not introducing new artifacts.

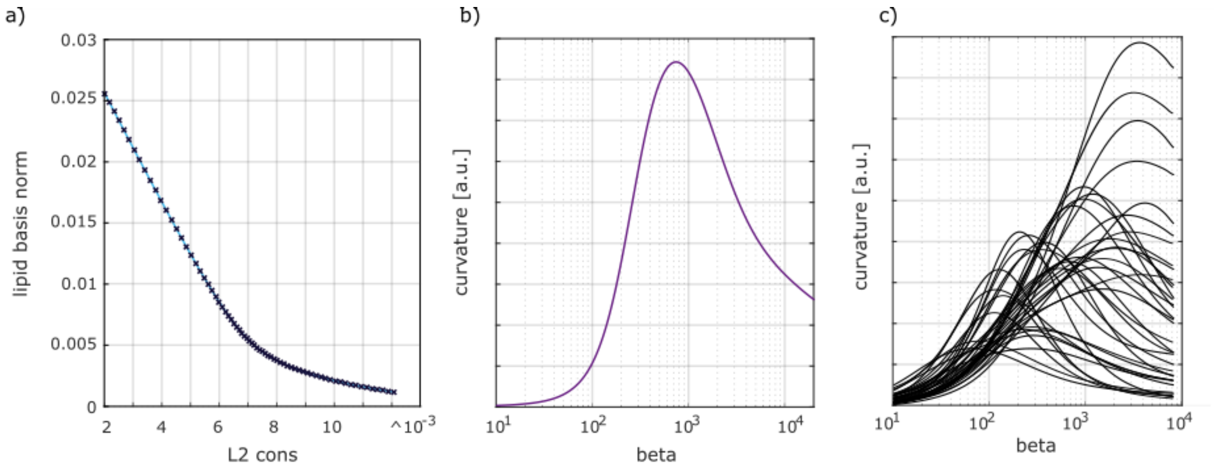


*Figure A: L-curve (lipid basis norm vs. L2 of consistency) and the corresponding curvature (b) from one sample slice. Curvatures from different slices and volunteers (c).*

5. Cabanes E, Confort-Gouny S, Le Fur Y, Simond G, Cozzone PJ. Optimization of Residual Water

Signal Removal by HLSVD on Simulated Short Echo Time Proton MR Spectra of the Human Brain. *J Magn Reson*. 2001;150(2):116-125. doi:10.1006/jmre.2001.2318

6. Kay SM. *Modern Spectral Estimation: Theroy and Application*.; 1988. doi:978-0130151599

7. Bilgic B, Chatnuntawech I, Fan AP, et al. Fast image reconstruction with L2-regularization. *J Magn Reson Imaging*. 2014;40(1):181-191. doi:10.1002/jmri.24365

8. Hansen PC. The L-Curve and its Use in the Numerical Treatment of Inverse Problems. *Comput Inverse Probl Electrocardiol*. 2000;(4):119-142. doi:10.1.1.33.6040

**LCModel .control File:**

```
$LCMODL
LTABLE=0
LCSV=11
LPS= 8
LCOORD= 9
hzpppm= 399.719
deltat= 1.2500e-04
nunfil= 1036
neach= 50
nratio = 0
ppmend= 1.8
ppmst= 4.2
doecc= F
degzer=0
sddegp= 1
sddegz= 2
dows= T
rfwhm = 1.45
wconc = 55510
dkntmn= 0.25
nsimul = 0
ndslic= 1
ndrows= 48
ndcols= 48
islice= 1
irowst= Irow
irowen= Irow
icolst= Jcol
icolen= Jcol
shifmx(2)= 0.01
shifmn(2)= -0.01
$END
```

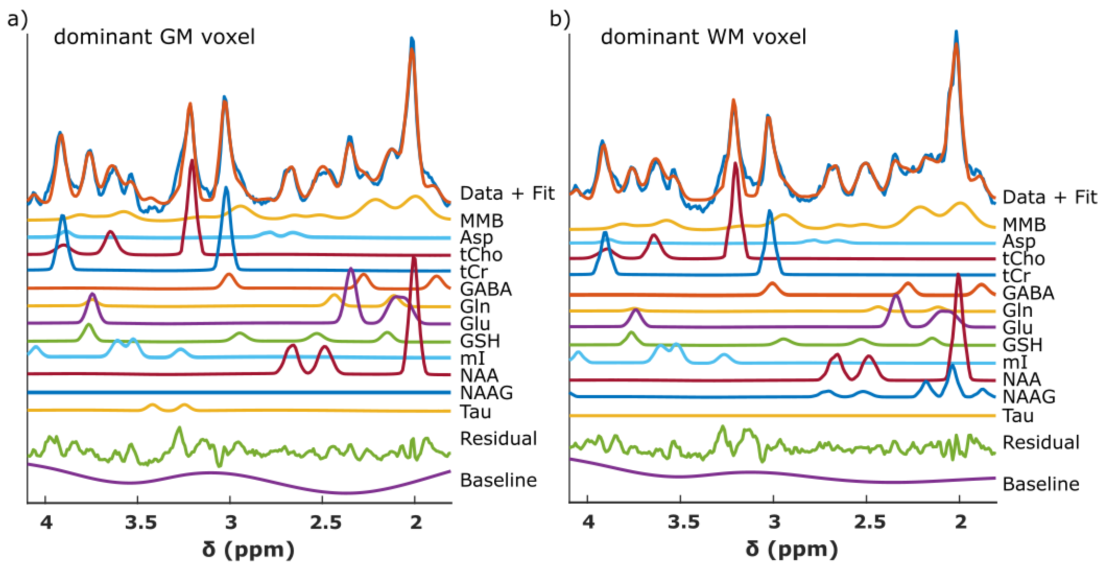


*Figure B: Sample spectra from dominant gray matter (a) and white matter (b) voxel with all fitted metabolites for the configuration L2-MM-0.25.*

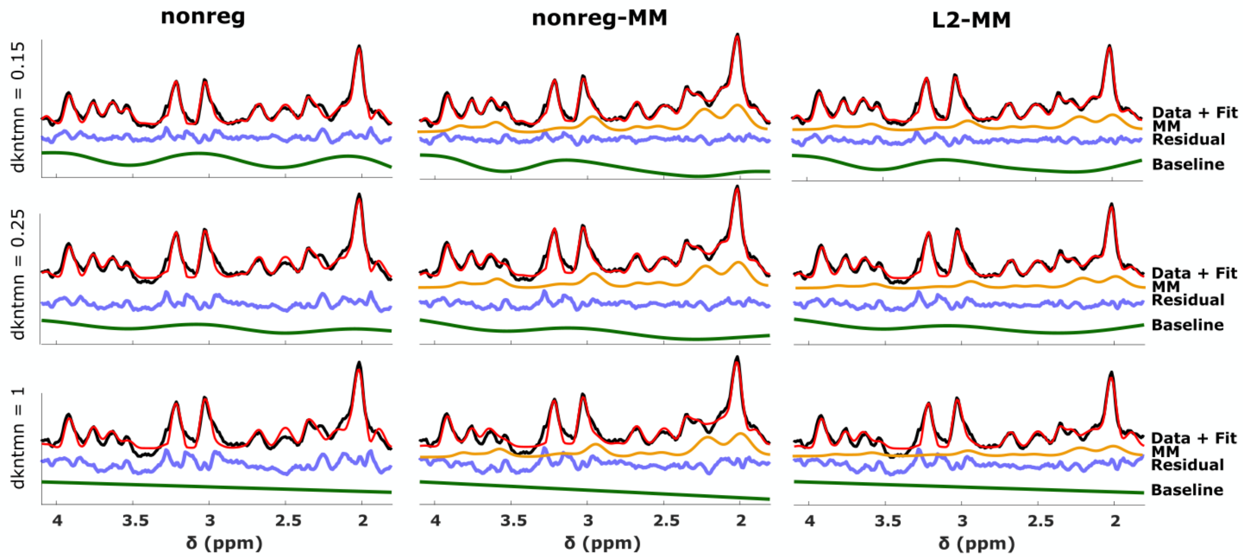


*Figure C: Data, fit, MM spectrum, residual and spline baseline for each configuration from one sample spectrum of volunteer V5.*

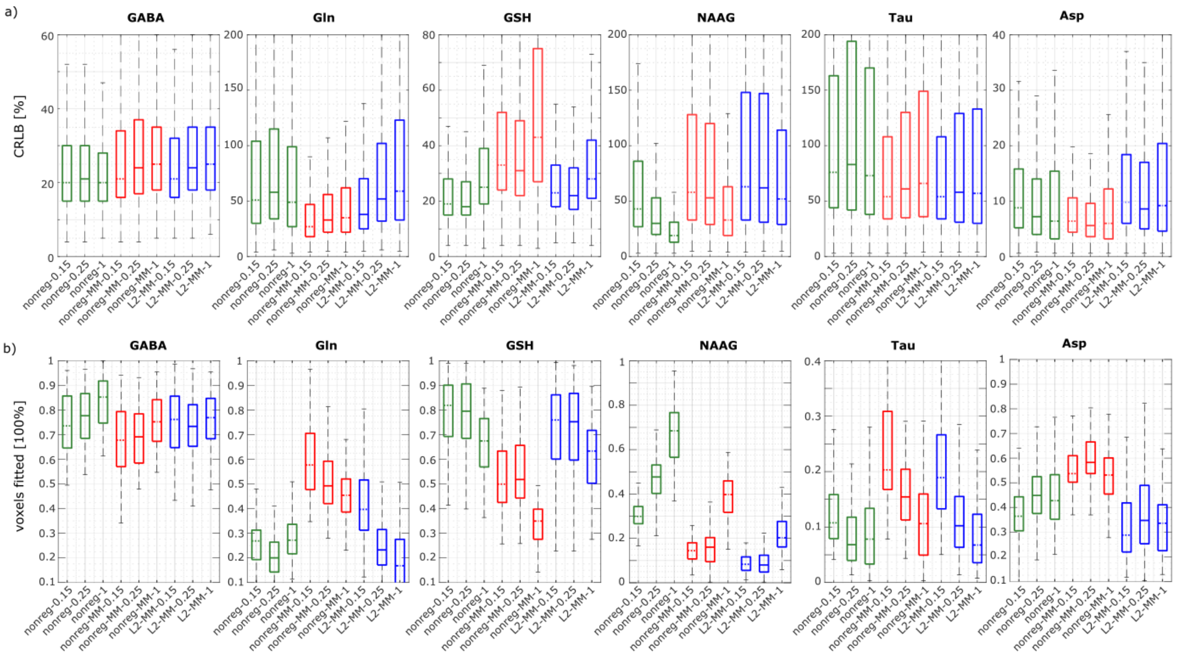


*Figure D: CRLB (a) and voxels fitted (b) from LCModel for all configurations and different metabolites. Data from all measurements, volunteers and slices are pooled.*

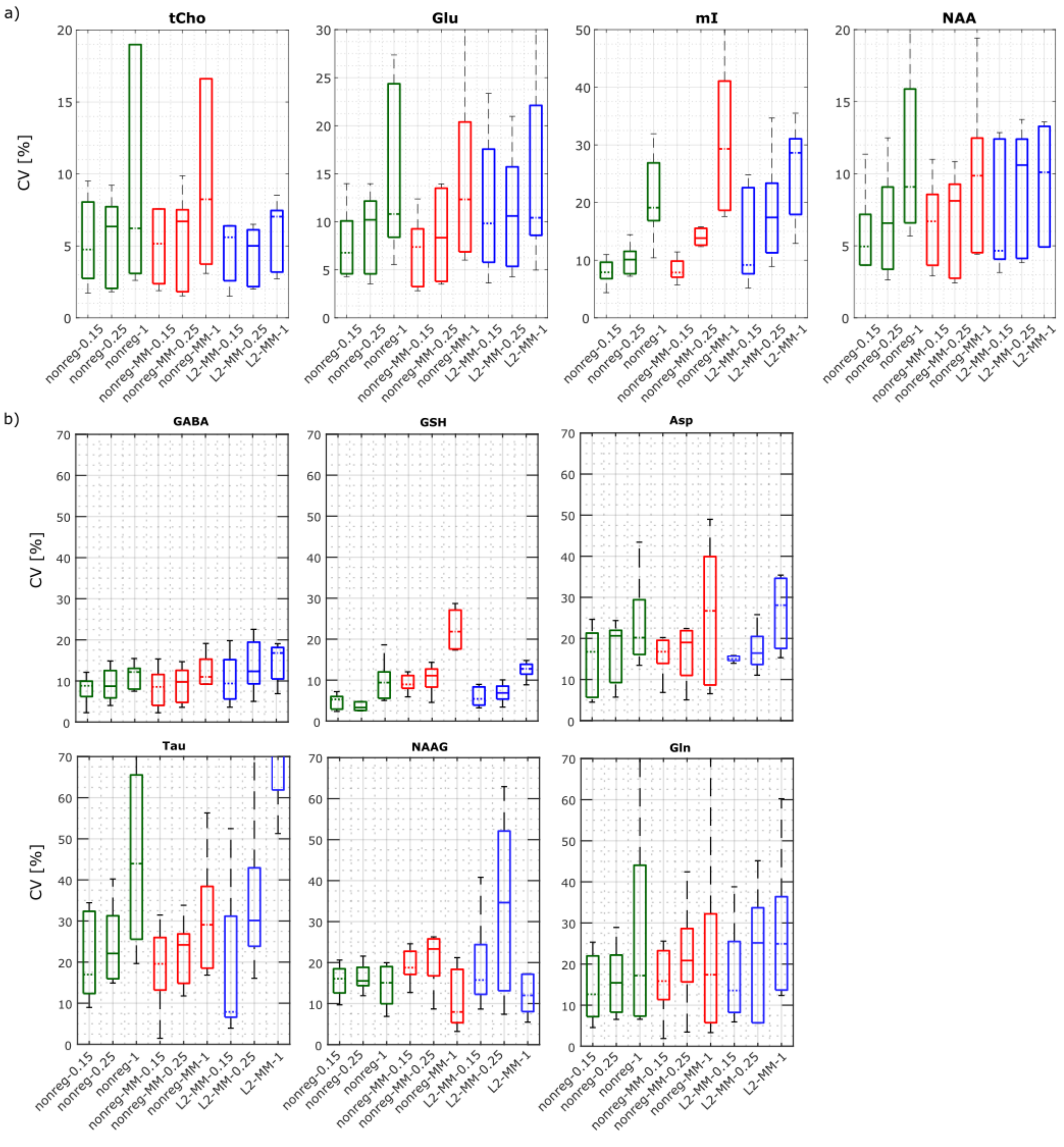


*Figure E: Coefficient of variance of the metabolite ratios (/tCr) for the test-retest measurements for all configurations and ten metabolite concentration ratios. Data pooled from all volunteers and a) all slices, b) slice 6-9.*

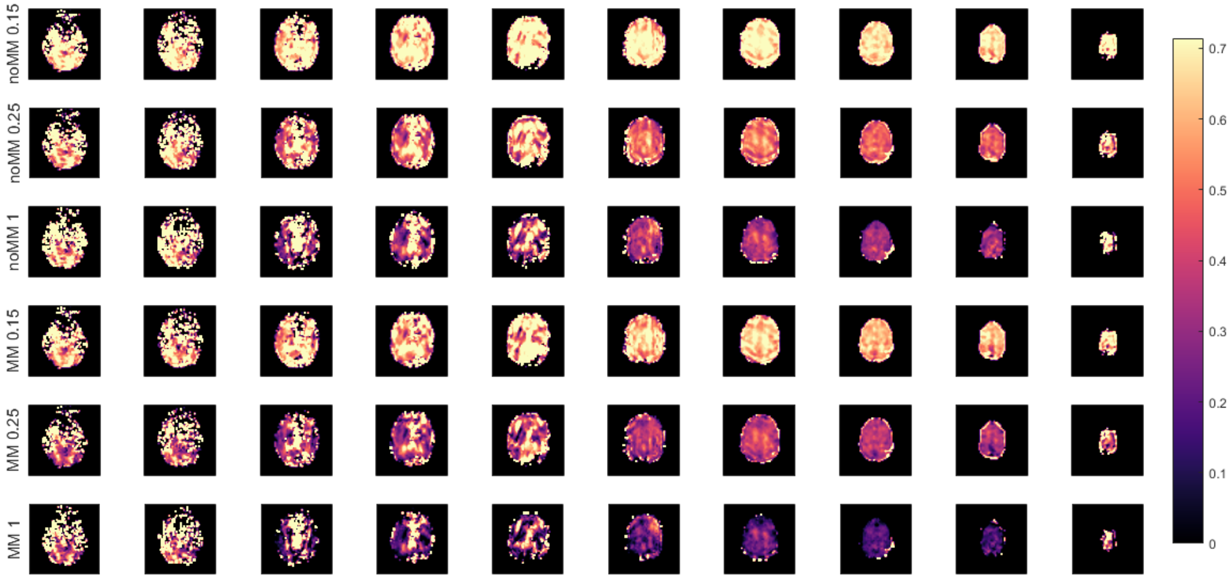


*Figure F: mI/tCr concentration ratio maps for the configurations: nonreg-0.15, nonreg-0.25, nonreg-1 and nonreg-MM-0.15, nonreg-MM-0.25, nonreg-MM-1 for volunteer 5. Data reported in arbitrary units.*

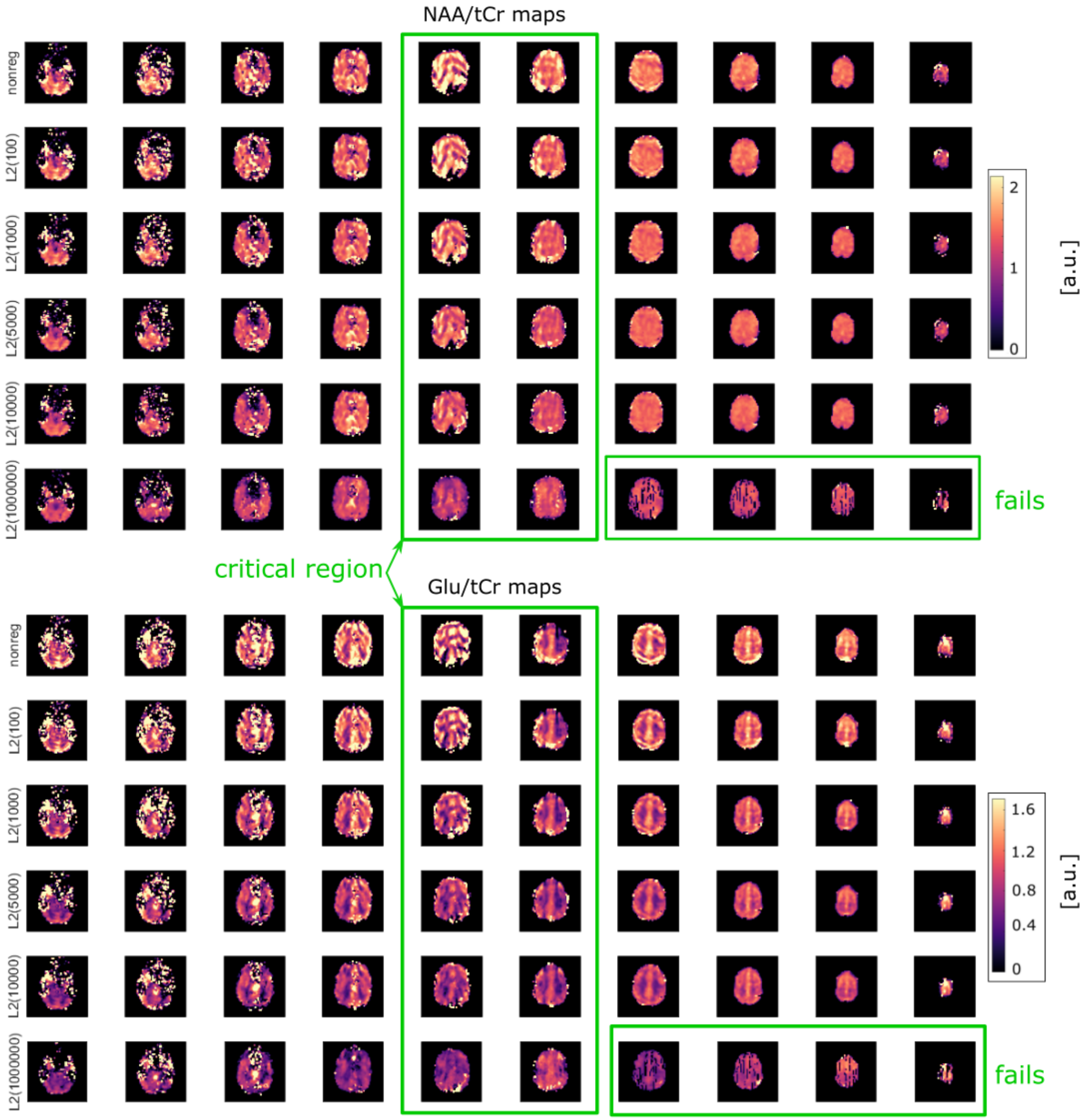


*Figure G: Comparison of metabolite ratio maps (NAA/tCr and Glu/tCr) for different regularization parameters: First row shows non-regularized data, the rows below show L2-regularized data for β=100, 1000, 5000, 10000 and $10^6$, respectively.*

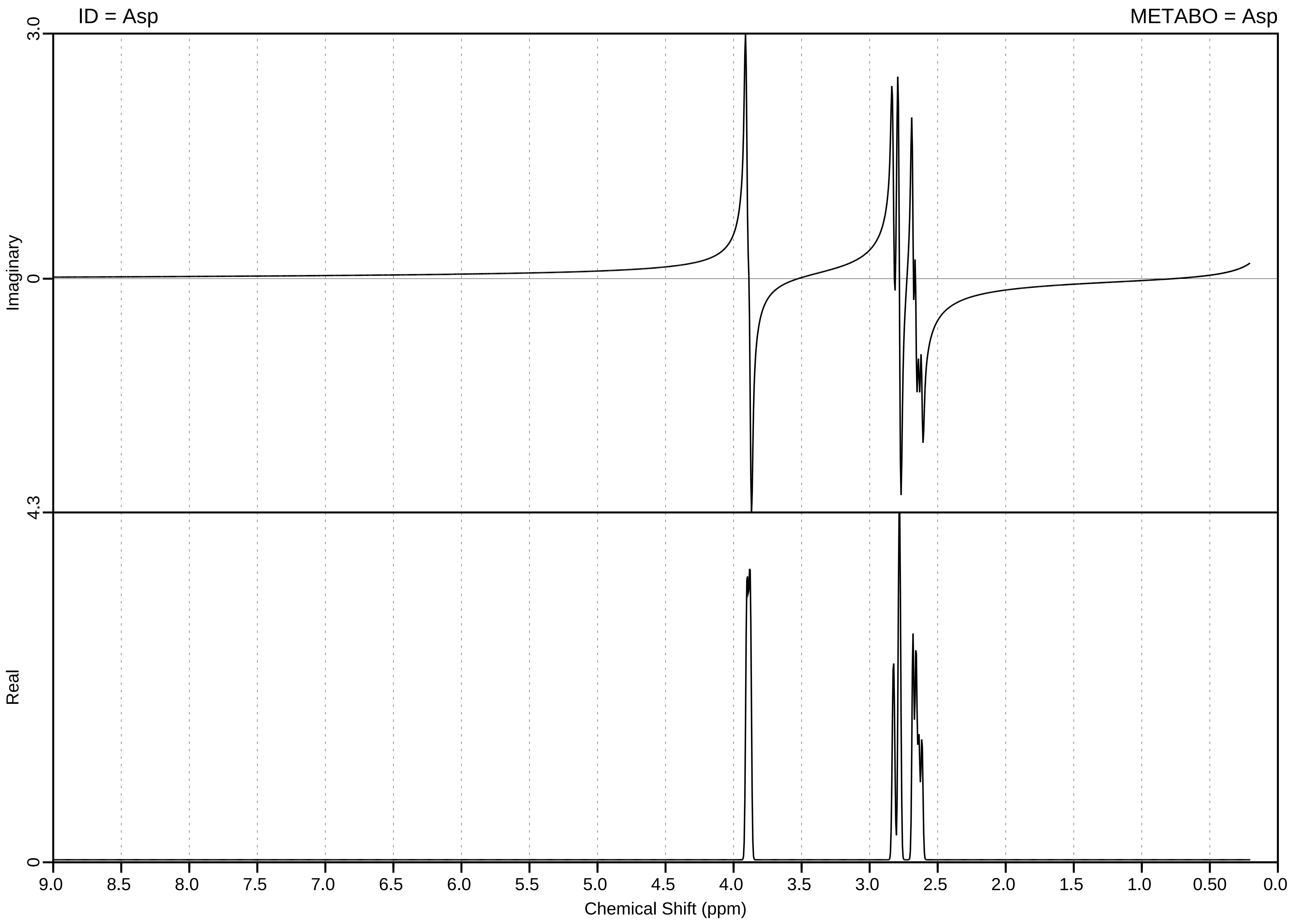

ID = Asp
METABO = Asp
Imaginary
Real
3.0
0
4.3
0
9.0
8.5
8.0
7.5
7.0
6.5
6.0
5.5
5.0
4.5
4.0
3.5
3.0
2.5
2.0
1.5
1.0
0.50
0.0
Chemical Shift (ppm)

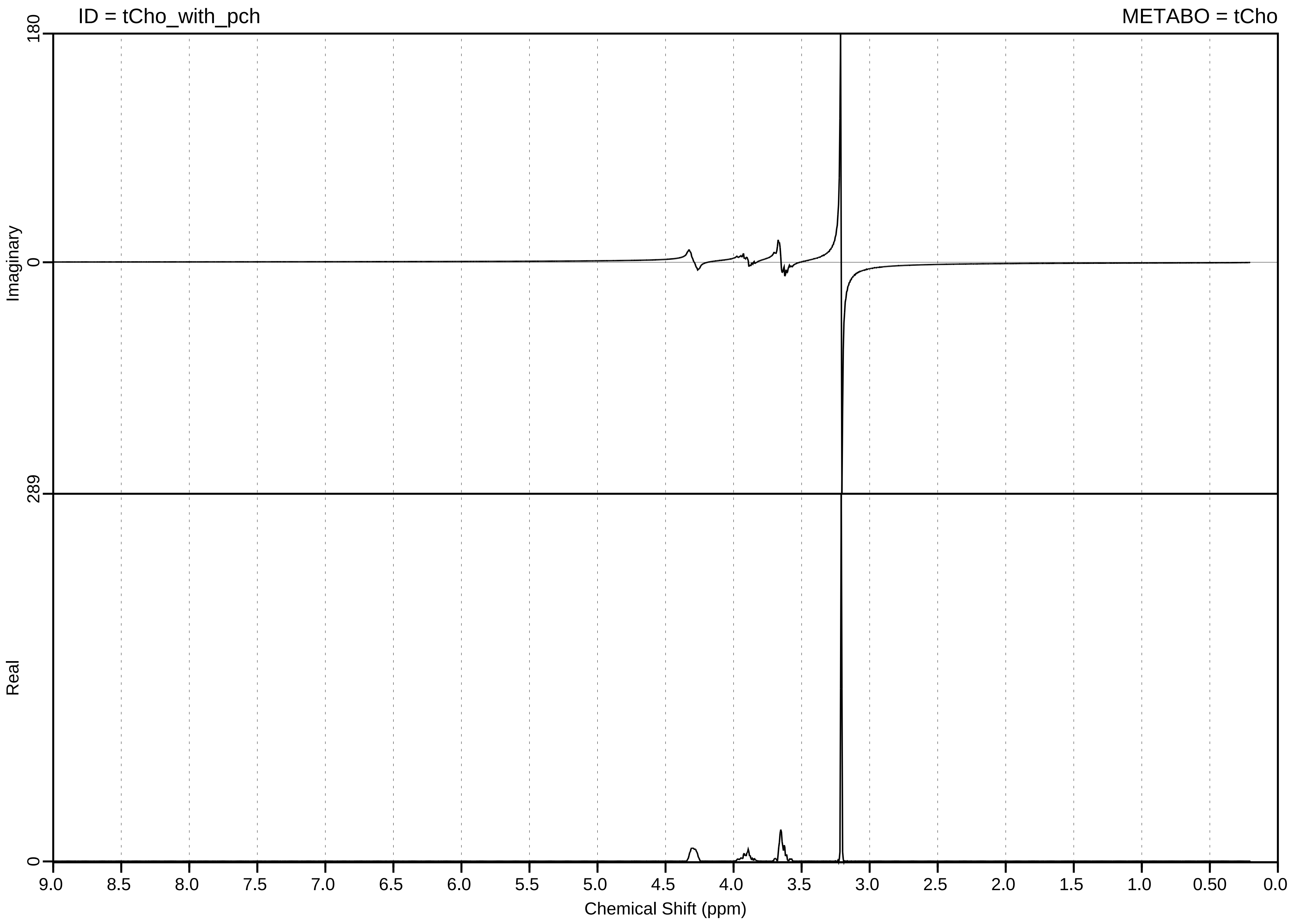

ID = tCho_with_pch
METABO = tCho
Imaginary
Real
180
0
289
0
9.0
8.5
8.0
7.5
7.0
6.5
6.0
5.5
5.0
4.5
4.0
3.5
3.0
2.5
2.0
1.5
1.0
0.50
0.0
Chemical Shift (ppm)

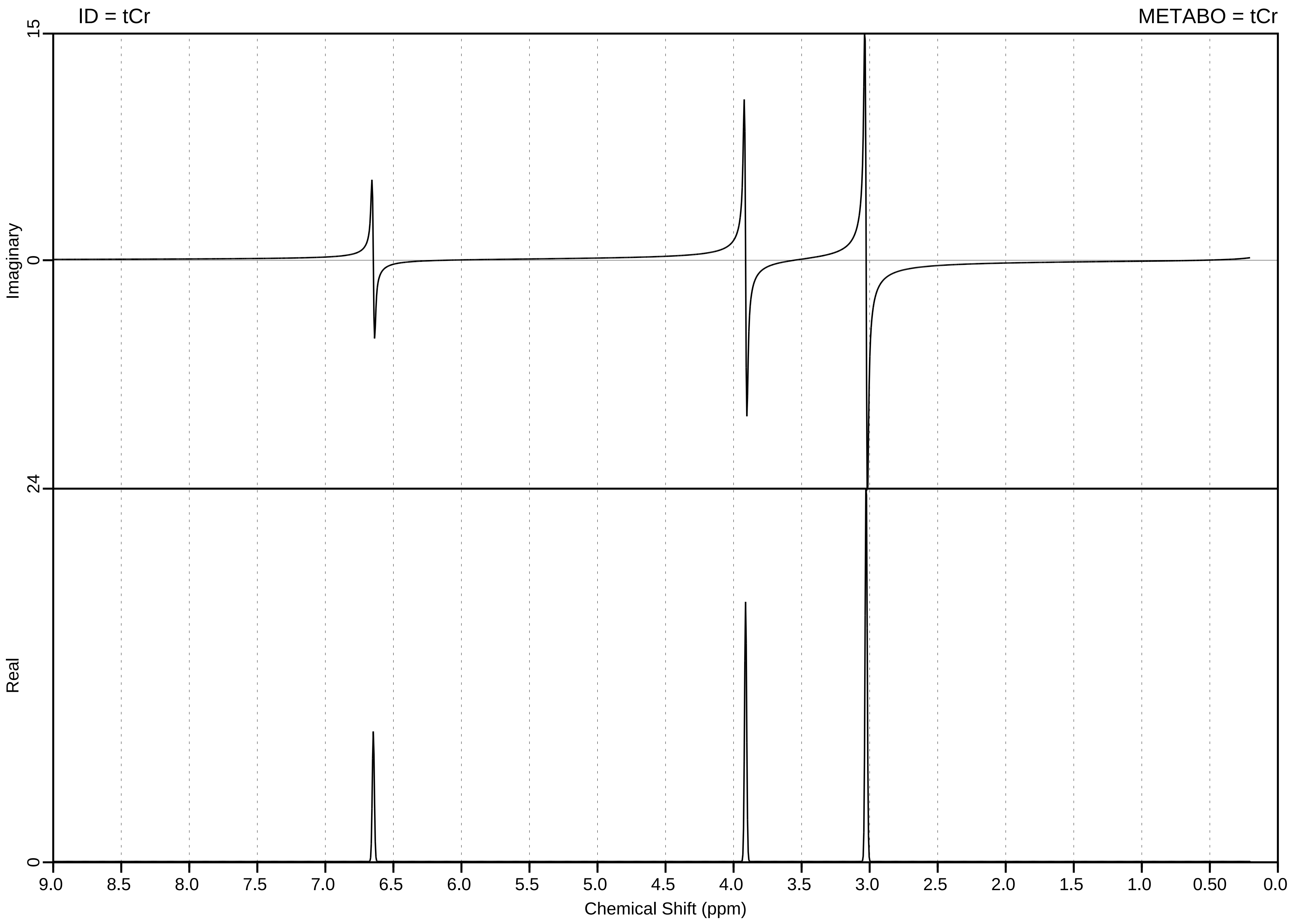

ID = tCr
METABO = tCr
15
Imaginary
0
24
Real
0
9.0
8.5
8.0
7.5
7.0
6.5
6.0
5.5
5.0
4.5
4.0
3.5
3.0
2.5
2.0
1.5
1.0
0.50
0.0
Chemical Shift (ppm)

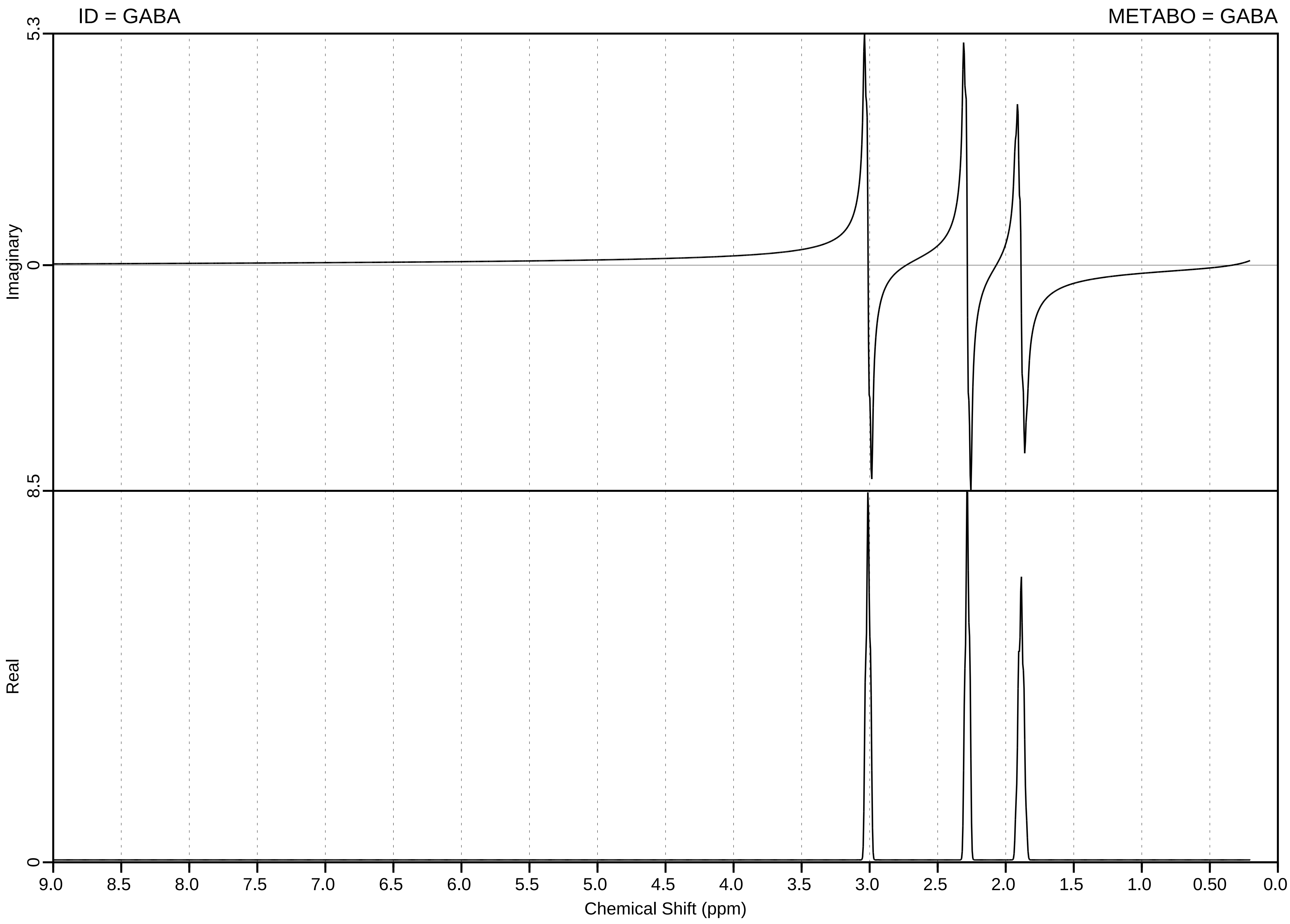

ID = GABA
METABO = GABA
5.3
0
8.5
0
Imaginary
Real
9.0
8.5
8.0
7.5
7.0
6.5
6.0
5.5
5.0
4.5
4.0
3.5
3.0
2.5
2.0
1.5
1.0
0.50
0.0
Chemical Shift (ppm)

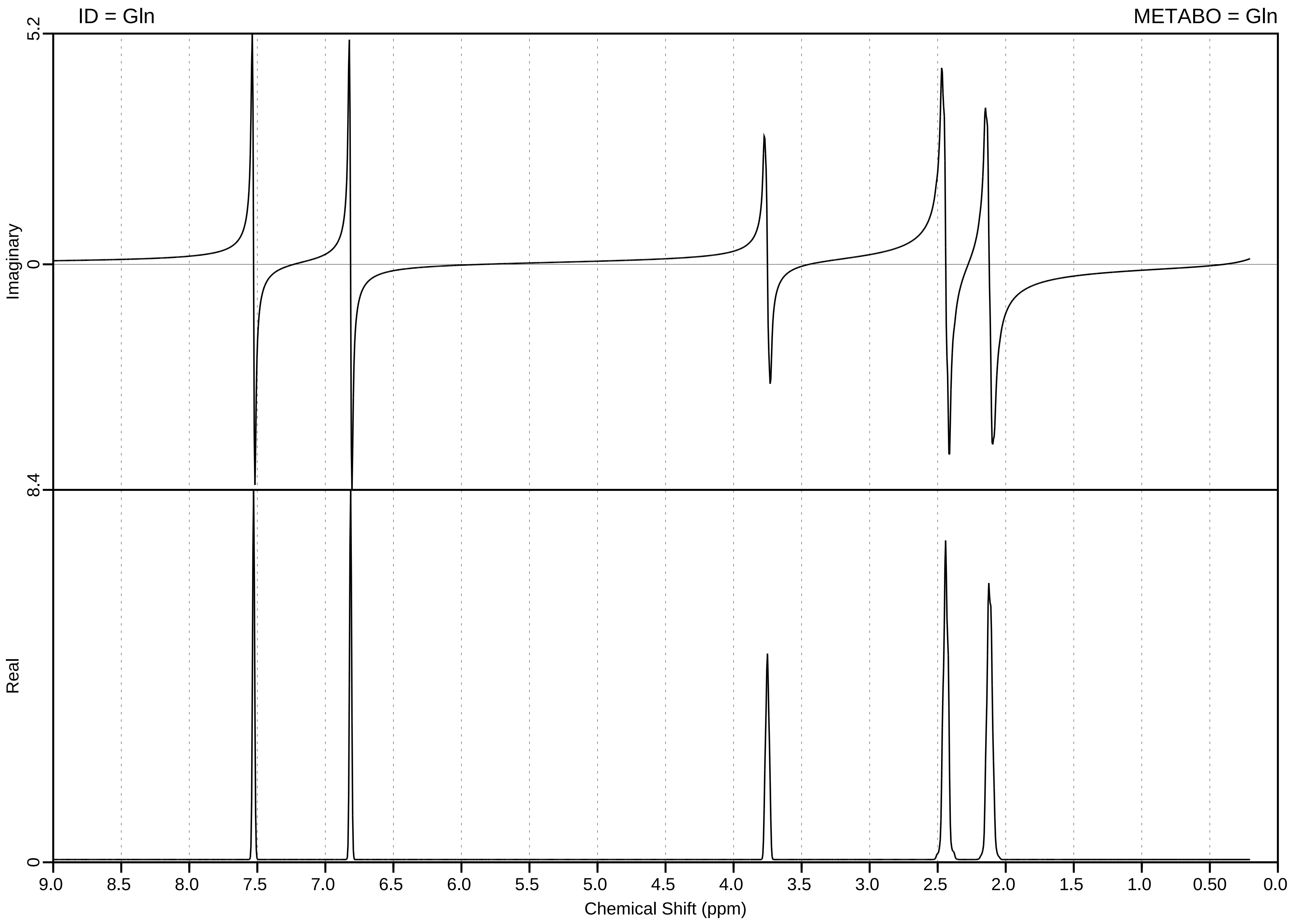
ID = Gln
METABO = Gln
Imaginary
Real
5.2
0
8.4
0
9.0
8.5
8.0
7.5
7.0
6.5
6.0
5.5
5.0
4.5
4.0
3.5
3.0
2.5
2.0
1.5
1.0
0.50
0.0
Chemical Shift (ppm)

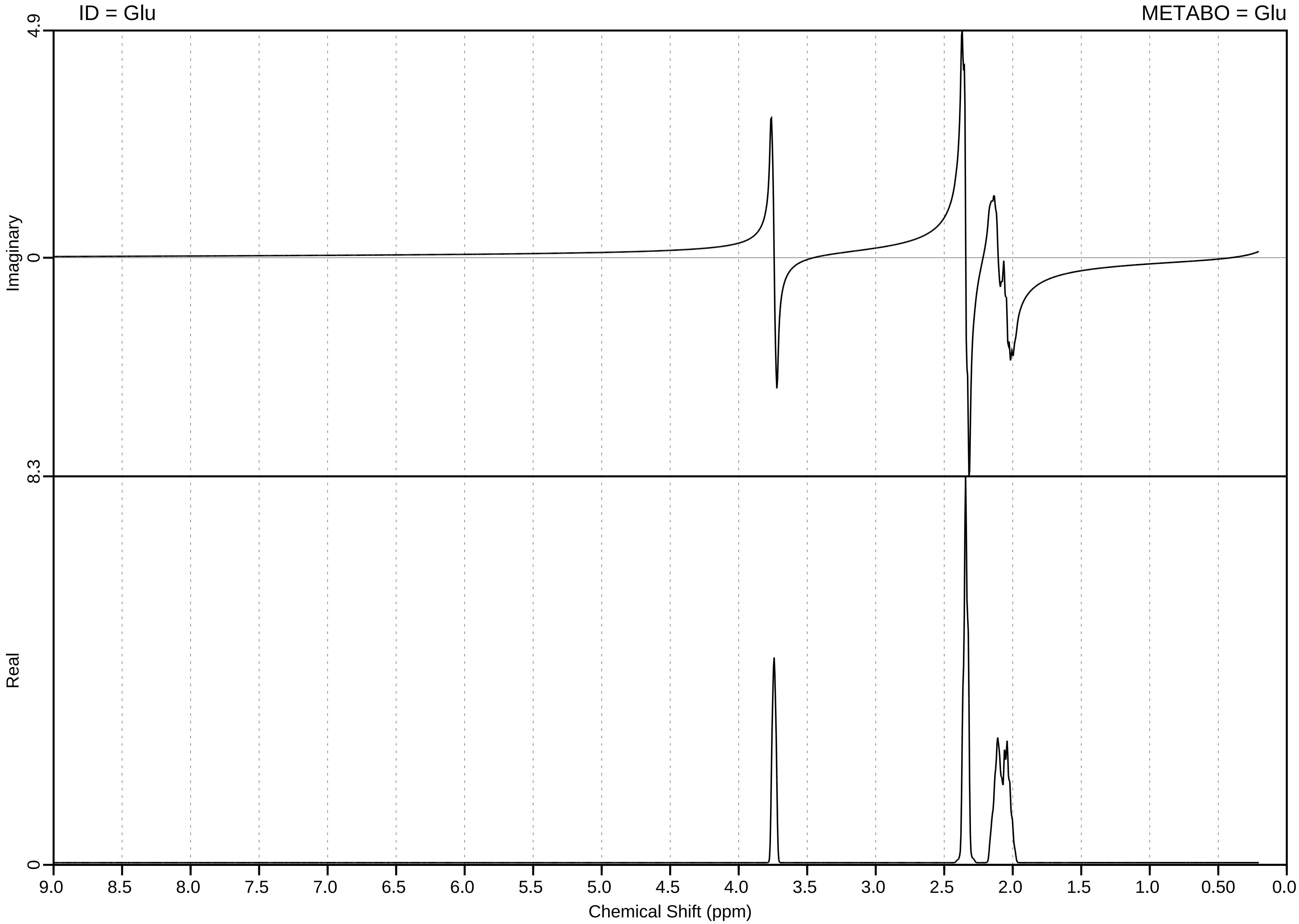

ID = Glu
METABO = Glu
Imaginary
Real
4.9
0
8.3
0
9.0
8.5
8.0
7.5
7.0
6.5
6.0
5.5
5.0
4.5
4.0
3.5
3.0
2.5
2.0
1.5
1.0
0.50
0.0
Chemical Shift (ppm)

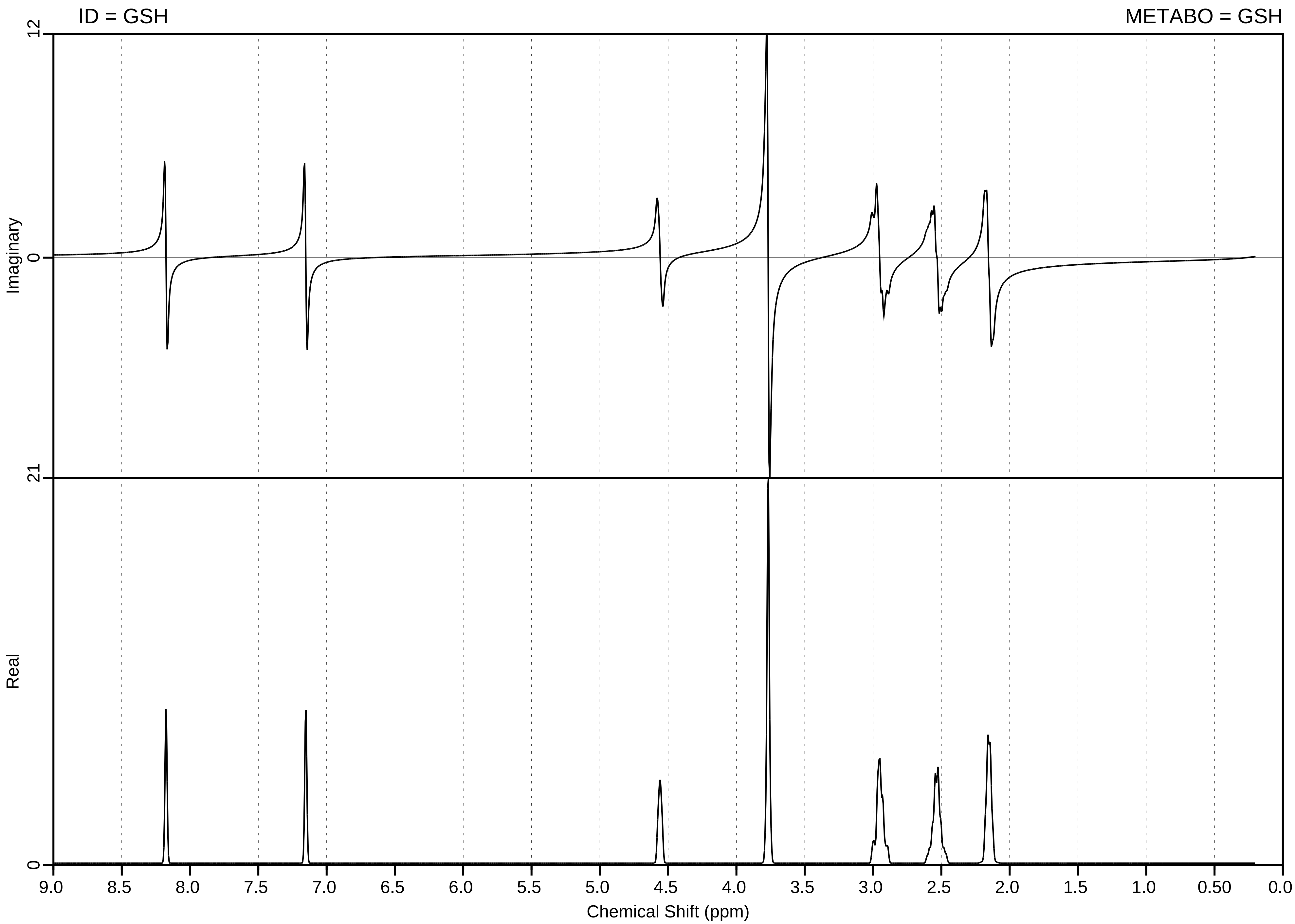

ID = GSH
METABO = GSH
Imaginary
Real
12
0
21
0
9.0
8.5
8.0
7.5
7.0
6.5
6.0
5.5
5.0
4.5
4.0
3.5
3.0
2.5
2.0
1.5
1.0
0.50
0.0
Chemical Shift (ppm)

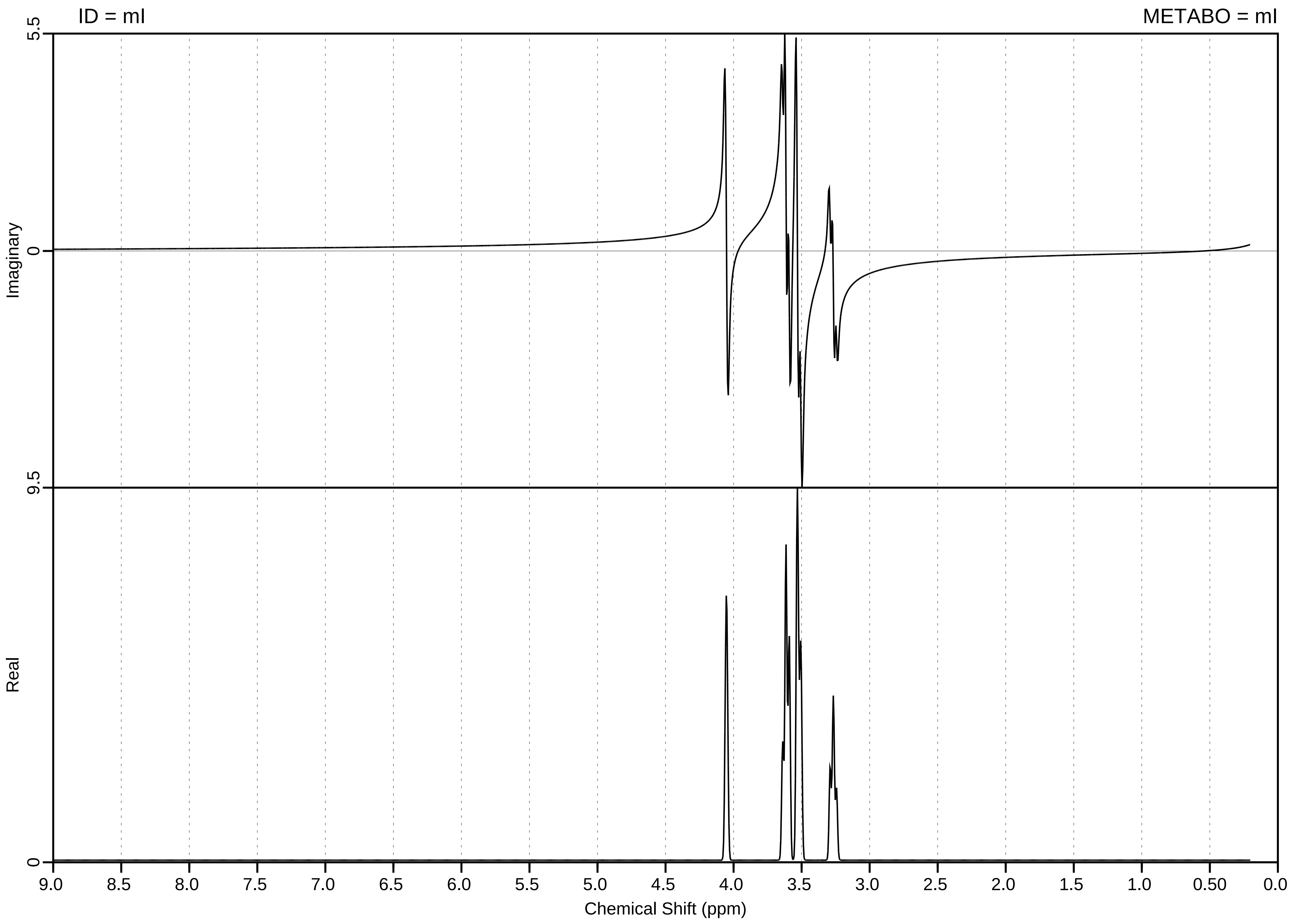

ID = mI
METABO = mI
5.5
0
Imaginary
9.5
0
Real
9.0
8.5
8.0
7.5
7.0
6.5
6.0
5.5
5.0
4.5
4.0
3.5
3.0
2.5
2.0
1.5
1.0
0.50
0.0
Chemical Shift (ppm)

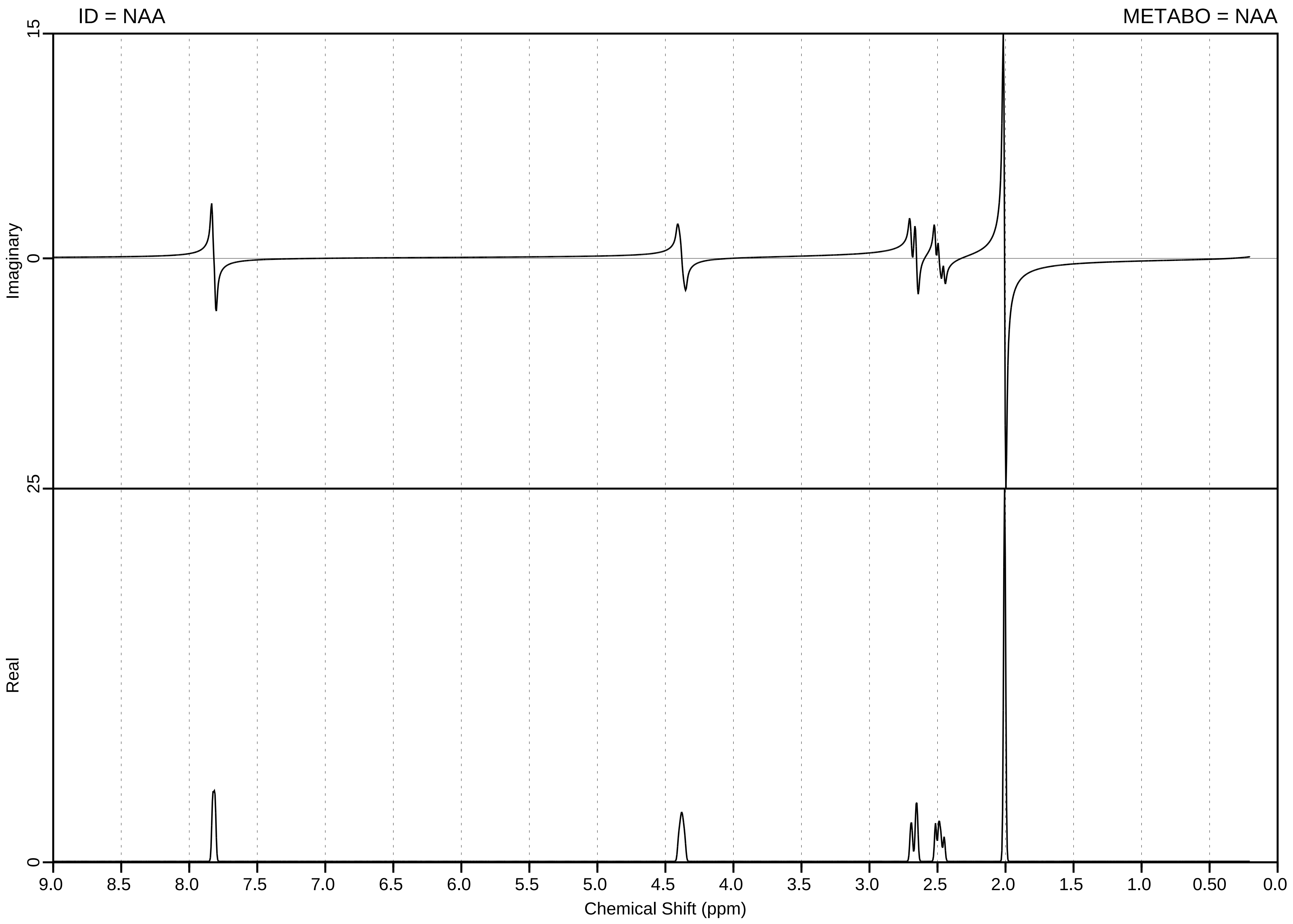

ID = NAA
METABO = NAA
Imaginary
Real
15
0
25
0
9.0
8.5
8.0
7.5
7.0
6.5
6.0
5.5
5.0
4.5
4.0
3.5
3.0
2.5
2.0
1.5
1.0
0.50
0.0
Chemical Shift (ppm)

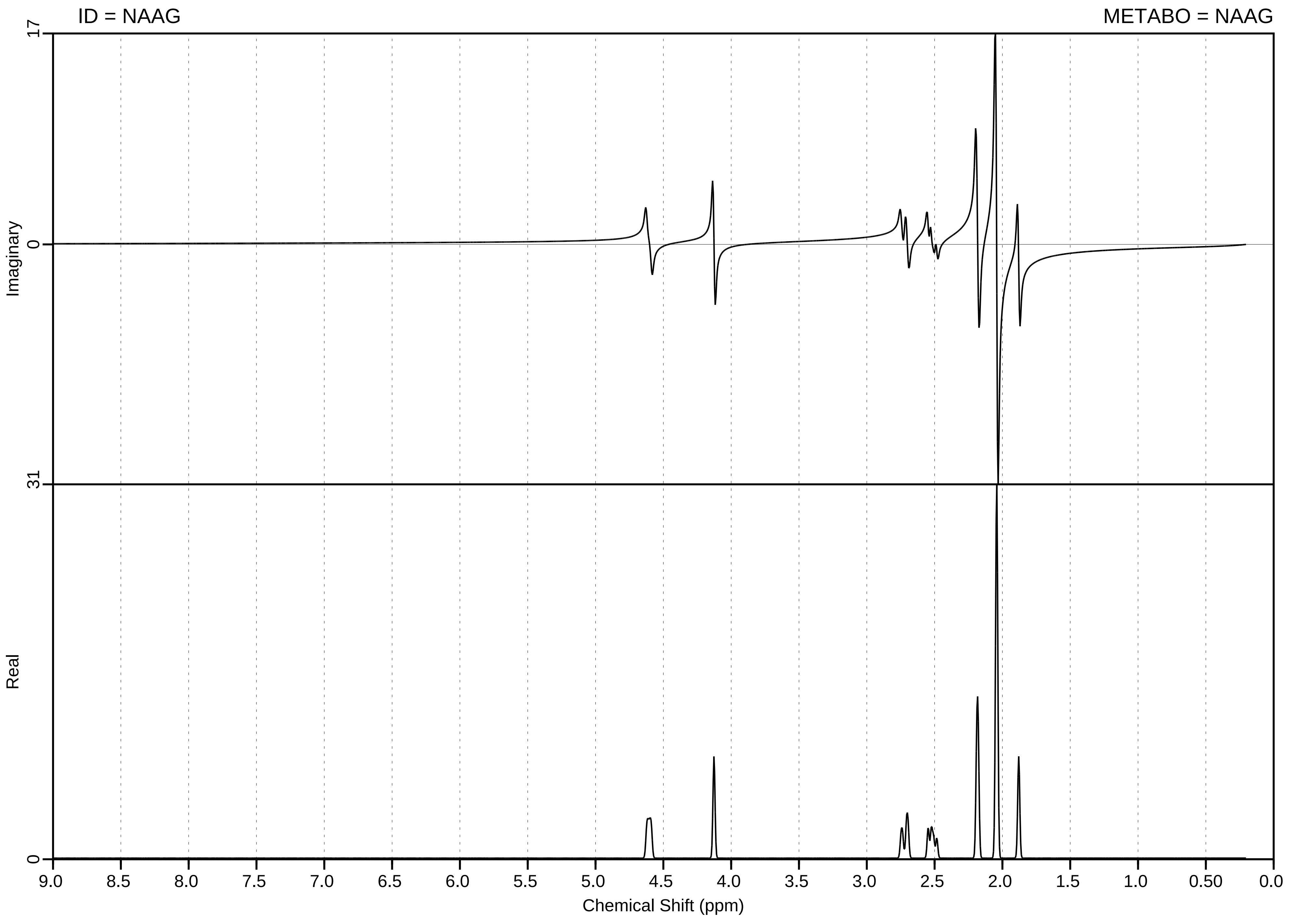

ID = NAAG
METABO = NAAG
17
0
31
0
Imaginary
Real
9.0
8.5
8.0
7.5
7.0
6.5
6.0
5.5
5.0
4.5
4.0
3.5
3.0
2.5
2.0
1.5
1.0
0.50
0.0
Chemical Shift (ppm)

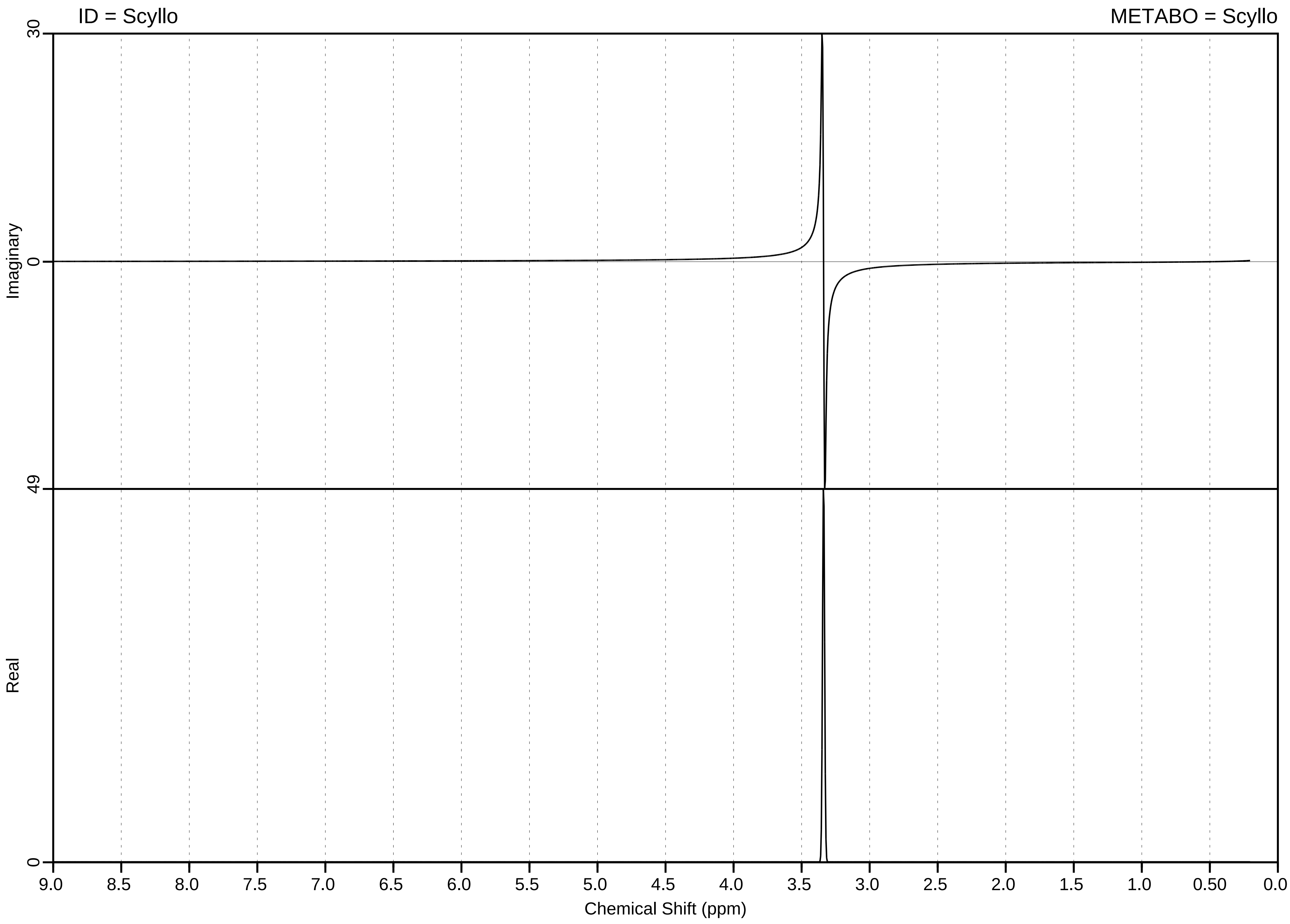
ID = Scyllo
METABO = Scyllo
Imaginary
Real
30
0
49
0
9.0
8.5
8.0
7.5
7.0
6.5
6.0
5.5
5.0
4.5
4.0
3.5
3.0
2.5
2.0
1.5
1.0
0.50
0.0
Chemical Shift (ppm)

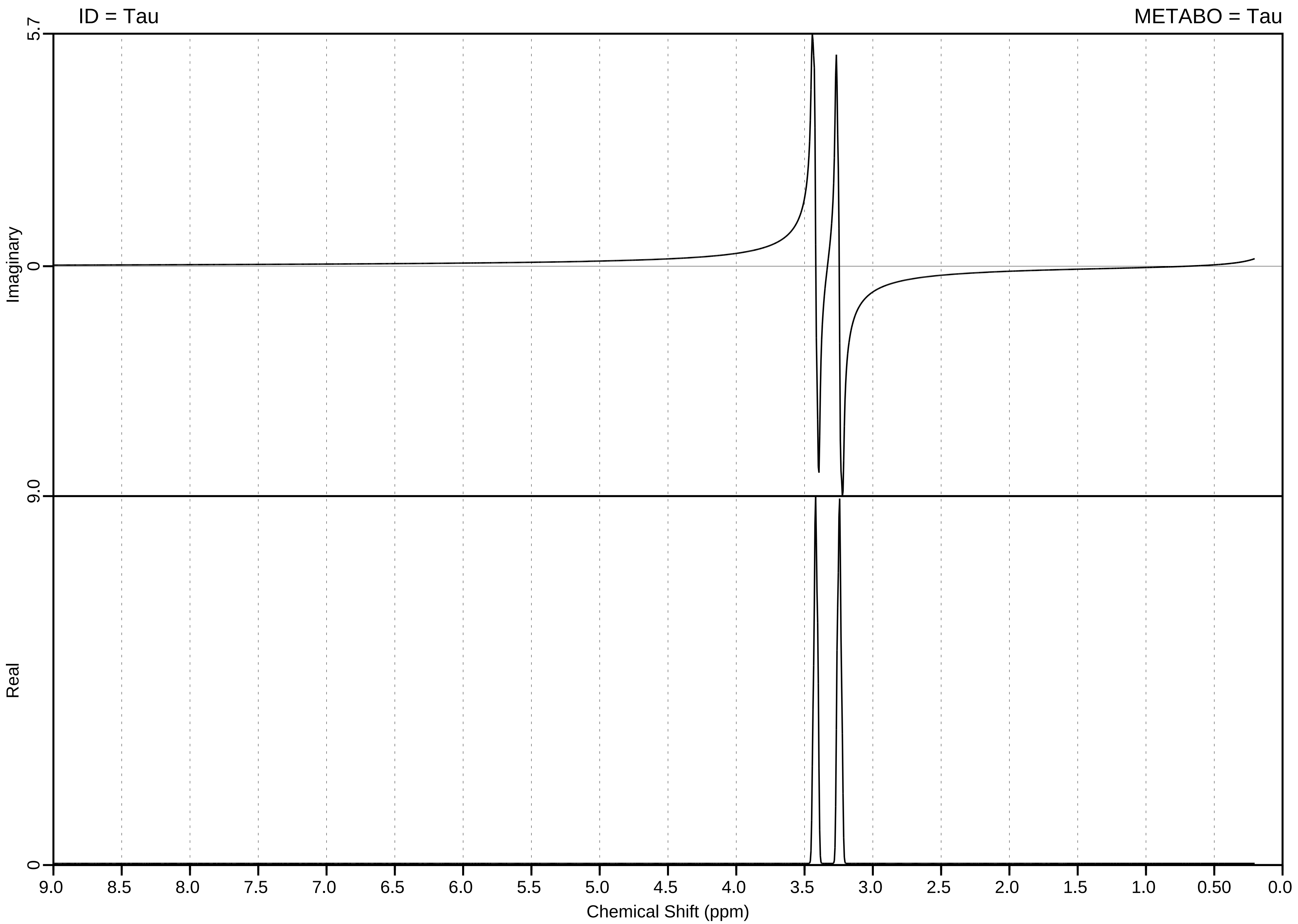

ID = Tau
METABO = Tau
Imaginary
Real
5.7
0
9.0
0
9.0
8.5
8.0
7.5
7.0
6.5
6.0
5.5
5.0
4.5
4.0
3.5
3.0
2.5
2.0
1.5
1.0
0.50
0.0
Chemical Shift (ppm)

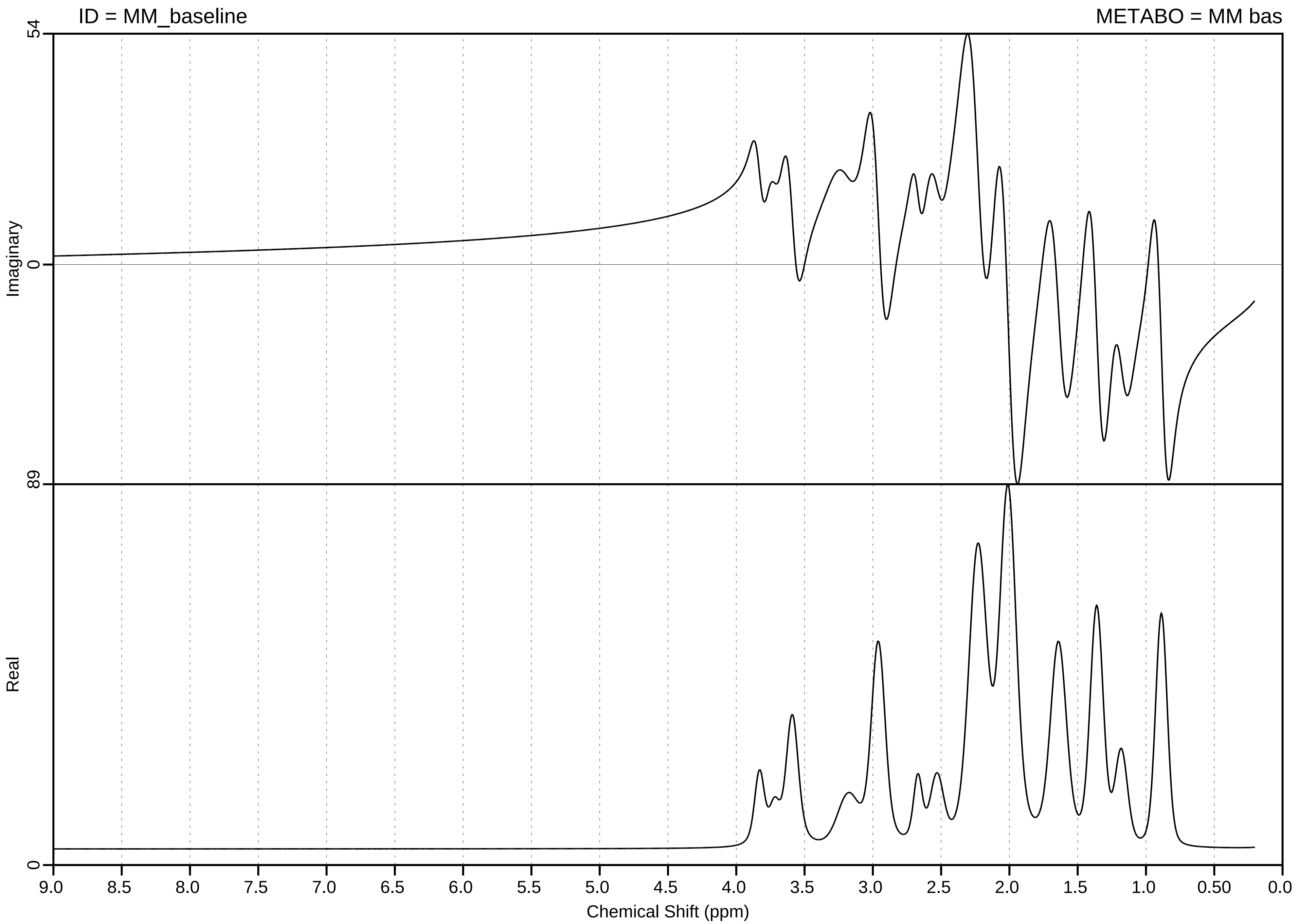

ID = MM_baseline
METABO = MM bas
54
Imaginary
0
89
Real
0
9.0
8.5
8.0
7.5
7.0
6.5
6.0
5.5
5.0
4.5
4.0
3.5
3.0
2.5
2.0
1.5
1.0
0.50
0.0
Chemical Shift (ppm)
